# Long non-coding RNA Neat1 regulates adaptive behavioural response to stress in mice

**DOI:** 10.1101/773333

**Authors:** Michail S. Kukharsky, Natalia N. Ninkina, Haiyan An, Vsevolod Telezhkin, Wenbin Wei, Camille Rabesahala de Meritens, Johnathan Cooper-Knock, Shinichi Nakagawa, Tetsuro Hirose, Vladimir L. Buchman, Tatyana A. Shelkovnikova

**Author notes:** To whom correspondence should be addressed at Medicines Discovery Institute, Cardiff University, Main Building, Cardiff, CF10 3AT, United Kingdom.

## Abstract

NEAT1 is a highly and ubiquitously expressed long non-coding RNA (lncRNA) which serves as an important regulator of cellular stress response. However, the physiological role of NEAT1 in the central nervous system (CNS) is still poorly understood. In the current study, we addressed this by characterising the CNS function in the Neat1 knockout mouse model (*Neat1^-/-^* mice), using a combination of behavioural phenotyping, electrophysiology and expression analysis. RNAscope® *in situ* hybridisation revealed that in wild-type mice, Neat1 is expressed evenly across the CNS, with high expression in glial cells and low expression in neurons. Loss of Neat1 in mice results in an inadequate reaction to physiological stress manifested as hyperlocomotion and panic escape response. In addition, *Neat1^-/-^* mice display deficits in social interaction and rhythmic patterns of activity but retain normal motor function and memory. *Neat1^-/-^* mice do not present with neuronal loss, overt neuroinflammation or gross synaptic dysfunction in the brain. However, cultured *Neat1^-/-^* neurons are characterised by hyperexcitability and dysregulated calcium homeostasis, and stress-induced neuronal activity is also augmented in *Neat1^-/-^* mice *in vivo*. Gene expression analysis showed that Neat1 may act as a weak positive regulator of multiple genes in the brain. Furthermore, loss of Neat1 affects alternative splicing of genes important for the CNS function and implicated in neurological diseases. Overall, our data suggest that Neat1 is involved in stress signaling in the brain and fine-tunes the CNS functions to enable adaptive behaviour in response to physiological stress.

## Introduction

Long noncoding RNAs (lncRNAs) are an arbitrary group comprising non-protein coding transcripts longer than 200 nucleotides. The majority of lncRNAs display tissue-, cell- and context-specific expression, and only few of them are ubiquitous and highly expressed. The prominent member of the latter group is Nuclear Paraspeckle Assembly Transcript 1 (NEAT1), a nuclear-retained lncRNA with important roles in cellular (patho)physiology and one of the most intensely studied lncRNAs. NEAT1 contributes to various cellular processes, from regulation of transcription and chromatin active state to miRNA biogenesis (reviewed in Ref. (Fox et al., 2017)). *NEAT1* locus produces two transcripts, NEAT1_1 and NEAT1_2. The longer NEAT1 isoform, NEAT1_2, is essential for the assembly of nuclear bodies termed paraspeckles (Sunwoo et al., 2009; Fox and Lamond, 2010), whereas NEAT1_1, albeit also a paraspeckle component, is dispensable for their formation and likely plays various paraspeckle-independent roles (Barry et al., 2017b). NEAT1 expression is elevated in stressed cells, such as those subjected to hypoxia (Choudhry et al., 2015), viral infection (Imamura et al., 2014), heat shock (Lellahi et al., 2018), mitochondrial stress (Wang et al., 2018) or proteasome inhibition (Hirose et al., 2014).

Altered *NEAT1* gene expression has been connected to various pathological conditions. Changes in NEAT1 levels is a recurrent theme in neoplasias, and the gene is a hotspot for mutations in several types of cancer (Fox et al., 2017). According to the Genotype-Tissue Expression (GTEx) database, *NEAT1* is expressed ubiquitously in the human body, with the lowest expression in the CNS, as compared to other organs and tissues (fig S1). However, altered *NEAT1* expression has been reported in all major neurodegenerative and psychiatric diseases, including frontotemporal dementia (FTD), Alzheimer’s, Huntington’s and Parkinson’s diseases, amyotrophic lateral sclerosis (ALS), epilepsy, traumatic brain injury and schizophrenia (reviewed in Ref. (An et al., 2018)). Mechanisms of NEAT1 transcripts involvement in the neurological conditions are still poorly understood, primarily because our knowledge of their function(s) is the CNS is still scarce. It is known *NEAT1* expression in neurons is responsive to neuronal activity (Barry et al., 2017a). Moreover, NEAT1 is able to modulate neuronal excitability, where its acute downregulation renders neurons more excitable (Barry et al., 2017a). NEAT1 transcripts regulate epigenetic marks on histones in neurons (Chakravarty et al., 2014; Butler et al., 2019). Because levels of NEAT1_2 in the intact brain are almost negligible (Nakagawa et al., 2011), NEAT1_1 can be considered as the main functional NEAT1 transcript in the CNS tissue under basal conditions. Despite the above findings, we still lack a clear picture of what and how NEAT1 contributes to neuronal function, especially at the organismal level.

*Neat1* knockout mouse strain was generated in 2011 through disruption of the *Neat1* promoter sequences, and these mice appeared to be viable and superficially normal (Nakagawa et al., 2011). However, subsequent more detailed studies revealed hormone dysfunction and decreased fertility of *Neat1* knockout females (Nakagawa et al., 2014), confirming an important role for Neat1 transcripts in specific physiological processes. *Neat1* knockout mice do not present with an overt neurological phenotype, however, neuronal deficits in these mice may only manifest under certain conditions such as the physiological stress experienced during pregnancy.

In the current study, we interrogated Neat1 function in the mammalian CNS using this mouse line. We show that loss of Neat1 perturbs normal behavioural responses of mice specifically under conditions of stress. This phenotype is not due to neuronal loss, neuroinflammation or gross changes in synaptic functions but rather can be attributed to altered neuronal excitability and changes in the alternative splicing of genes important for the CNS function. Our data suggest that Neat1 fine-tunes the CNS function under stressful conditions at the organismal level, which is consistent with the current view of NEAT1 as stress-responsive transcripts at the cellular level.

## Materials and methods

### Mouse colonies and genotyping

Generation of *Neat1^-/-^* mouse strain has been described previously (Nakagawa et al., 2011). The strain was maintained on the C57Bl/6J genetic background by backcrossing with wild-type mice from Charles River, UK. Primers used for mouse genotyping by PCR were 5’-CTAGTGGTGGGGAGGCAGT-3’ and 5’-AGCAGGGATAGCCTGGTCTT-3’.

Homozygous (*Neat1^-/-^*) and wild-type (*Neat1^+/+^*) mice were obtained by intercrossing hemizygous (*Neat1^+/-^*) mice. Experimental animals were housed individually, at 12 h light/12 h dark cycle, with food and water supplied *ad libitum*. Before behavioural testing, the animals were placed in the testing room in their home cages for 1 h for habituation. All work on animals was carried out in accordance with the United Kingdom Animals (Scientific Procedures) Act (1986).

### Behavioural testing

#### Locomotor activity

Spontaneous locomotor activity was assessed in the home-like cage (HC) equipped with infrared beams to monitor horizontal movements over a period of 36 h, under the 12 h light/12 h dark cycle. Number of beam breaks was recorded at 15-min intervals. The same level of illumination (as in the room used for normal housing (15-20 lux) was maintained during the day phase of the test. Locomotor activity under bright light conditions (600 lux) was recorded for 1 h using Ugo Basile 47420 multiple activity cage.

#### Elevated plus maze test

The test was performed on a standard plus-shaped apparatus. The test was carried out under bright light (400 lux). Each mouse was placed in the central square of the maze facing one of the open arms and was allowed to freely explore the maze for 5 min. Video recordings were analysed to determine the duration and frequency of entries into open and closed arms, number of head dips and stretch-attend postures (Walf and Frye, 2007; Komada et al., 2008).

#### Y-maze test

Working memory was assessed using a Y-maze apparatus with arms measuring 40×6×20 cm (LxWxH). The test was conducted under dim light conditions (15-20 lux). Spontaneous alternations were assessed in the two-trial test (Dellu et al., 2000; Hughes, 2004). Briefly, mice were initially allowed to explore two arms of maze for 5 min with the third arm blocked. After a 30 min inter-trial interval, the second trial was conducted, where mice were allowed to explore all three arms for 5 min. An arm entry was recorded when more than half of the mouse body crossed the border between two arms. Number of entries and time spent in each arm were registered.

#### Resident-intruder test

To assess aggression and social interaction, the standard protocol of resident-intruder test was used (Winslow, 2003; Koolhaas et al., 2013). Animals were maintained without bedding changed for 1 week before the task. The cage top was removed and the male intruder was placed in the cage for 5 min. The number of attacks and total time spent in a physical interaction were recorded.

#### Social odour test

Social odours were collected from a home cage where five C57Bl/6 adult male mice had been housed for 6 days without bedding change. The bottom of the cage was wiped with a cotton swab in a zig-zag pattern to collect social odour. Mice were left to habituate in a new cage in the presence of an unused cotton swab for 30 min prior to test. Subsequently, mice were exposed to a new cotton swab without odour, followed by a cotton swab with a social odour. The amount of time spent sniffing each swab over a period of 2 min was recorded.

#### Restraint stress

To induce *cFos* expression, mice were exposed to a 60 min restraint stress. Animals were closed into custom-made cylindrical polycarbonate tubes with ventilation holes measuring 12.7 cm long and 3.8 cm in diameter, which allowed to significantly restrict their movements. Control animals were left undisturbed in their home cages in the same procedure room. After the restraint, mice were returned to their home cages for 60 min. Control and experimental mice were euthanised immediately after, and brain tissue was fixed for analysis as described below.

### RNA isolation and gene expression analysis

Total RNA was extracted using PureLink total RNA extraction kit (Life Technologies) and possible DNA contamination was removed using RNase free DNase kit (Qiagen). cDNA synthesis was performed on 500 ng of total RNA using SuperScript IV reverse transcriptase (Life Technologies) and random hexamer primers (Promega). Quantitative real-time PCR was run in triplicate on an StepOne™ real-time PCR instrument and data were analyzed using StepOne™ Software v2.0 (all Applied Biosystems, Life Technologies). Gapdh was used for normalisation. Primer seqeunces were as follows: Neat1 total 5’-TGGAGATTGAAGGCGCAAGT3’ and 5’-ACCACAGAAGAGGAAGCACG-3’; Neat1_2 5’-AACTACCAGCAATTCCGCCA-3’ and 5’-GAGCTCGCCAGGTTTACAGT-3’; Malat1 5’-GAGCTCGCCAGGTTTACAGT-3’ and 5’-AACTACCAGCAATTCCGCCA-3’; Gapdh 5’-AAGAGGGATGCTGCCCTTAC-3’ and 5’-TACGGCCAAATCCGTTCACA-3’; Gabrr2 5’-ACATCCAAGCCAAGCCATTTG-3’ and 5’-CATGGTGAAGTCGTGGTCCT-3’; Gabra1 5’-TATGGACAGCCCTCCCAAGA-3’ and 5’-TACACGCTCTCCCAAACCTG-3’; Gabra2 5’-ACAGATTCAAAGCCACTGGAGG-3’ and 5’-TCTTCTTGTTGCCAAAGCTG-3’; Rfk 5’-GGCCACATTGAGAATTTCCCC-3’ and 5’-GGTGGTGAGCATTGGATGGA-3’.

### RNA sequencing and analysis

Total RNA was prepared as described above and its quality examined using Agilent 4200 TapeStation. RNA sequencing was performed at the School of Biosciences Genomics Research Hub. Libraries were prepared using the TruSeq stranded mRNA kit (Illumina). Paired-end sequencing was carried out on Illumina NextSeq500 (read length: 75 bp; coverage ∼50 million reads/sample). Reads were aligned to the mouse reference genome (GRCm38) using STAR (Dobin et al., 2013), and Fragments Per Kilobase of transcript per Million mapped reads (FPKM) values were obtained using DESeq2 (Love et al., 2014). Sequencing reads were viewed in the IGV browser (Thorvaldsdottir et al., 2013). Ensembl IDs for genes identified by DESeq2 were mapped to known genes and tested for GO term enrichment using DAVID 6.8 online tool (https://david.ncifcrf.gov/). Differential splicing events were identified using the Leafcutter pipeline (Li et al., 2018b). GO term and pathway enrichment analysis of differentially spliced genes was performed using the Enrichr online tool (https://amp.pharm.mssm.edu/Enrichr/).

### Western blotting

SDS-PAGE loading buffer was used for direct homogenisation of tissue, followed by denaturation at 100°C for 5 min. After SDS-PAGE, proteins were transferred to PVDF or nitrocellulose membrane (GE Healthcare) by semi-dry blotting. Membranes were blocked in 4% milk in TBST, incubated in primary antibody at 4⁰ C overnight and in HRP-conjugated secondary antibody (GE Healthcare) for 1.5 h at RT. For detection, WesternBright Sirius kit (Advansta) and Bio-Rad ChemiDoc™ Gel Imaging System were used. Equal loading was confirmed by re-probing membranes with an antibody against beta-actin.

### RNA in situ hybridisation and immunohistochemistry in mouse tissue

Mouse brains and spinal cords were fixed in 4% PFA overnight and embedded in paraffin wax. Eight µm thick sections were mounted on poly-L-lysine coated slides (ThermoScientific). For RNAscope® ISH analysis, Ms-Neat1-short (440351) probe (Advanced Cell Diagnostics), with the target region 1416 – 2381 in mouse Neat1, was used according to manufacturer’s instructions. For Nissl staining, sections were incubated in 0.5% Cresyl Violet Acetate solution (C5042, Sigma) and differentiated in acidified ethanol. Immunohistochemistry with 3,3’-diaminobenzidine (DAB) as a substrate was performed as described earlier (Peters et al., 2012). Images were taken using Leica DMRB microscope or Olympus BX40 slide scanner (x20 magnification). Quantification of neuron numbers was performed using the 3D Object Counter plugin of ImageJ software (https://imagej.nih.gov/ij/).

### Primary antibodies

Primary antibodies used for Western blot and immunohistochemistry: GFAP (rabbit polyclonal, G9269, Sigma); Iba1 (rabbit monoclonal, ab178846, Abcam); PSD95 (DLG4) (rabbit polyclonal, 20665-1-AP, Proteintech), synaptophysin (mouse monoclonal, 611880, BD Biosciences); riboflavin kinase (rabbit polyclonal, 15813-1-AP, Proteintech); GAD67 (rabbit polyclonal, 10408-1-AP, Proteintech); cFos (mouse monoclonal, 66590-1-Ig, Proteintech); beta-actin (mouse monoclonal A5441, Sigma).

### Primary mouse hippocampal cultures

Primary cultures of mouse hippocampal neurons were prepared from P1 litters of WT and Neat1^-/-^ mice as described previously (Kukharsky et al., 2015). Neurons were transfected with pEGFP-C1 plasmid using Lipofectamine2000 (Life Technologies) and imaged 24 h post-transfection, using BX61 microscope equipped with F-View II camera and CellF software. Day *in vitro* (DIV) 7 neurons were used for analysis. Neurite analysis was performed using “Sholl Analysis” and “Simple Neurite Tracer” plugins of ImageJ.

### Electrophysiological recordings and analysis

Patch-clamp - voltage and current recordings were made in DIV7 *Neat1^+/+^* and *Neat1^-/-^* neurons using conventional patch-clamp in the whole-cell configuration (Hamill et al., 1981) employing Axopatch 200B amplifier interfaced to a computer running pClamp 9 using a Digidata 1322A A/D interface (Molecular Devices). All electrophysiological studies were performed at a controlled room temperature of 22 ± 0.5 °C. Recordings were digitized at 10 kHz and low-pass filtered at 2 Hz using an 8-pole Bessel filter. The standard bath solution (pH 7.4) contained: 135mM NaCl, 5mM KCl, 1.2 MgCl2, 1.25 mM CaCl2, 10 mM D-glucose, 5 mM HEPES. The standard pipette solution (pH 7.2) contained: 117 mM KCl, 10 mM NaCl, 11 mM HEPES, 2 mM Na2-ATP, 2 mM Na-GTP, 1.2 mM Na2-phosphocreatine, 2 mM MgCl2, 1 mM CaCl2 and 11 mM EGTA. Mean resting membrane potential (Vm) of the neurons were determined during 120 s gap-free recording periods in current clamp mode (I = 0 pA). Spike analysis was performed on the first iAP using Clampfit 9. Na+ currents were recorded using a standard voltage-step protocol, as described elsewhere (Telezhkin et al., 2016).

### Ca^2+^ imaging

Imaging was performed using Olympus IX71 inverted microscope and fluorescence source (Xenon arc or LED), alongside with a rapid perfusion system. Neurons were loaded with Fura-2 AM (ThermoFisher, 4µM final concentration) in full media for 45 min under standard culturing conditions. Cells were transferred into basal recording solution for imaging (135 mM NaCl, 5 mM KCl, 2 mM CaCl2, 1 mM MgCl2, 10 mM glucose, 10 mM HEPES). Prior to KCl stimulation, basal recordings were taken for 60 sec. KCl (60 mM) solution was injected at this point, and recording continued for further 300 sec. ΔF/F0 was calculated as F(340 nm) / F(380 nm) and used to prepare the plot.

### Experimental Design and Statistical Analysis

To generate experimental cohorts, heterozygous *Neat1^-/-^*mice (C57Bl/6J background) were used. After genotyping of the progeny, animals were randomly assigned to the knockout (*Neat1^-/-^*) and WT control (*Neat1^+/+^*) groups. Evaluation of anxiety-like behavior, which can be affected by previous experience, was performed first, followed by tests for activity and memory. Assessment of social behavior, as the most stressful paradigm, was the last behavioural test conducted. Y-maze test was conducted in a procedure room other than rooms for locomotor activity tests, to exclude interaction with the surrounding cues. Resting time between tests was at least 7 days, and the order of tests was the same for all mice.

Statistical analysis was performed using GraphPad Prism 6 and STATISTICA 12 software. In all cases, results are presented as mean±SE, individual data are showed where appropriate, and *n* indicates the number of biological replicates. Effect size was calculated using Cohen’s d. Statistical details for each data set are indicated in the text and figure/table legends.

## Results

### Predominantly non-neuronal expression of Neat1 in the murine CNS

Neat1 levels in the murine organs and tissues, as studied by RNA *in situ* hybridisation, vary significantly, with high levels in the digestive tract, medium – in lung and kidney, and low – in skeletal muscle and brain (Nakagawa et al., 2011). However, the distribution of Neat1 within anatomical structures in the brain and its relative expression in different cell types in the CNS are still largely unknown. To address this, we used RNAscope® ISH with 5’ end specific Neat1 probe in the sagittal brain sections as well as in the spinal cord of adult (2-month old) wild-type (*Neat1^+/+^*) mice. As a negative control, we included brain sections from *Neat1* knockout (*Neat1^-/-^*) mice. This analysis revealed uniform expression of Neat1 throughout the murine brain and spinal cord, with low signal in neurons and substantial Neat1 accumulation in non-neuronal cells (the two cell types distinguished based on the size of nuclei) (Fig. 1A). Region-wise, the highest Neat1 level was detected in *corpus callosum*, *fornix* and *glia limitans*, which are the brain structures mainly composed of glial cells. In neuron-dense areas, such as hippocampus and granule cell layer of the cerebellum, Neat1 was much less abundant and present in a dotted pattern (Fig. 1A, insets - *hip* and *cer*, respectively). In contrast, in many non-neuronal cells, the entire nucleus was darkly stained, as was especially obvious in the *corpus callosum* (Fig. 1A, inset – *cc*). Neat1 ISH also resulted in the staining of the cytoplasmic perinuclear region in a population of cells expressing highest Neat1 levels, which we attributed to a staining artefact (Fig. 1A, *cc* - arrowheads). In *Neat1^-/-^* mice, only traces of Neat1 were detected in all anatomical brain regions except the cerebellum, where Neat1 signal was prominent in the granule cell layer (Fig. 1B). Similar to the brain, Neat1 signal was evenly distributed throughout the spinal cord of *Neat1^+/+^* mice, with low levels in neurons (Fig. 1C).

**Fig. 1.**
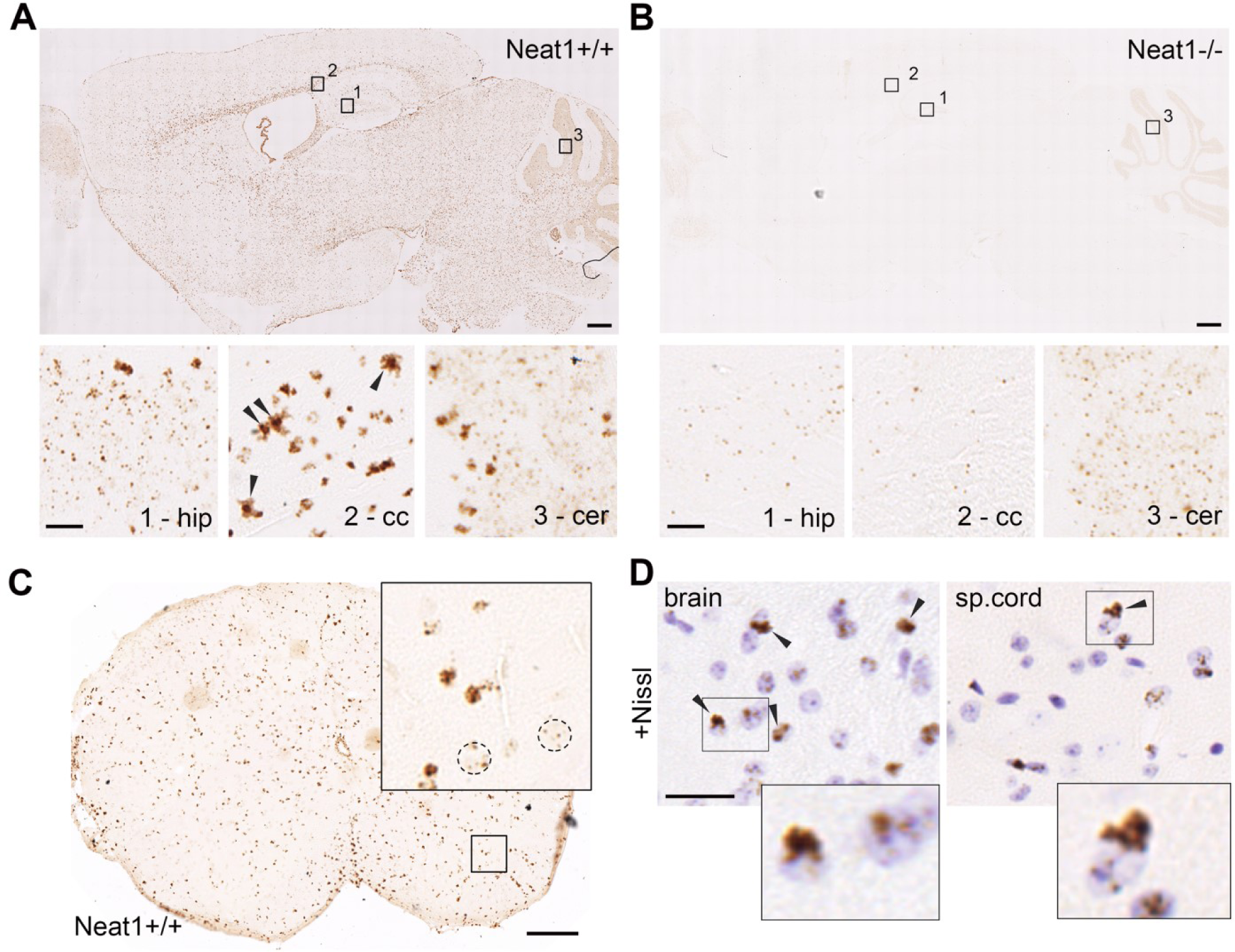
Neat1 is expressed in all cell types in the murine CNS. (**A,B**) Neat1 is expressed at high levels in non-neuronal cells and low levels in neurons in the murine brain. Neat1 i*n situ* hybridization (RNAScope® ISH, with 5’ fragment *Neat1* probe) on sagittal brain sections of *Neat1^+/+^* (**A**) and *Neat1^-/-^* (**B**) mice. Note low or absent Neat1 signal in all brain structures of *Neat1^-/-^* mice except the cerebellum (**B**). The bottom panels show insets from the top panels at higher magnification: 1 – granular layer of the hippocampus (dentate gyrus); 2 – *corpus callosum*; 3 – Purkinje cells and granular layer of the cerebellum. Arrowheads indicate cells expressing high levels of Neat1 with characteristic perinuclear (cytoplasmic) reactivity (presumably, ISH artefact). (**C**) Neat1 ISH in the *Neat1^+/+^*mouse spinal cord. Nuclei of motor neurons are circled. (**D**) Neat1 ISH with Nissl counterstain in the brain and spinal cord sections of a *Neat1^+/+^*mouse; arrowheads indicate satellite cells. Scale bars, (**A**, **B**) – 200 µm (main) and 20 µm (insets); (**C**) – 100 µm; (**D**) – 35 µm.

Counterstaining of brain and spinal cord sections with Nissl stain after the RNAscope® ISH allowed visualising large and lightly stained neuronal nuclei (where Neat1 signal was found as few dots per cell, Fig. 1D). Neat1 expression was highest in smaller nuclei often adjacent to neurons and corresponding to satellite cells – usually astrocytes and oligodendrocytes(Garcia-Cabezas et al., 2016) (Fig. 1D, arrowheads and inset). Of note, we were unable to achieve the typical Nissl staining pattern (i.e. staining of both nuclei and cytoplasm of neurons, as shown in Fig. 3B) after RNAscope® ISH, which could be due to the ISH processing protocol.

Therefore, Neat1 is expressed throughout the mammalian CNS, predominantly in glial cells (astrocytes and oligodendrocytes), and is present at low levels in neurons.

### Neat1 knockout mice demonstrate abnormal behavioural response to stress

Consistent with the original study (Naganuma et al., 2012), *Neat1^-/-^* mice showed no obvious deficits in general health, physical characteristics, or basic sensorimotor functions compared to their WT (*Neat1^+/+^*) littermates, except reduced body weight (Fig. 2A). However, we noticed that, despite normal behaviour while being observed in the cage without intervention, *Neat1^-/-^* mice exhibited a panic-like escape response and hyperlocomotion (excessive running and jumping) upon contact with an experimenter, e.g. handling or cage changes (Video S1). This phenotype gradually developed after weaning, and was apparent in 2-month old animals. We generated experimental groups of littermate *Neat1^-/-^* and *Neat1^+/+^*male mice (n=9 and 11, respectively) for behavioural analysis. During the period spanning from the age of 3 to 7 months, these mice were assessed using a panel of behavioral tests. First, we attempted to examine motor function in these cohorts using rotarod, however it was not possible due to multiple escape attempts of *Neat1^-/-^* mice after placing them on the apparatus. To assess the anxiety levels, we carried out the elevated plus maze (EPM) test. *Neat1^-/-^* mice presented with decreased anxiety-like behaviour in this test, as evidenced by a significantly higher number of entries to and increased time spent in open arms (Fig. 2B,C). However, an increase in the above readouts could be also interpreted as a behavioral phenotype of impulsivity (Ferguson and Gray, 2005; Pawlak et al., 2012) and/or high escape motivation (Holmes et al., 2000; Shoji et al., 2016). Consistent with the latter interpretation, the number of head dips from the open arm towards the floor was significantly increased, whilst the number of stretch-attend postures, when the rodent remains motionless with its body stretched forward, was decreased for *Neat1^-/-^* mice (Fig. 2D,E). This is indicative of escape attempts and inadequate risk assessment (Rodgers and Dalvi, 1997; Pawlak et al., 2012). There was no difference in the overall locomotor activity between the genotypes measured as the total number of arm entries (Fig. 2F), suggesting normal motor function in *Neat1^-/-^* mice.

**Figure 2.**
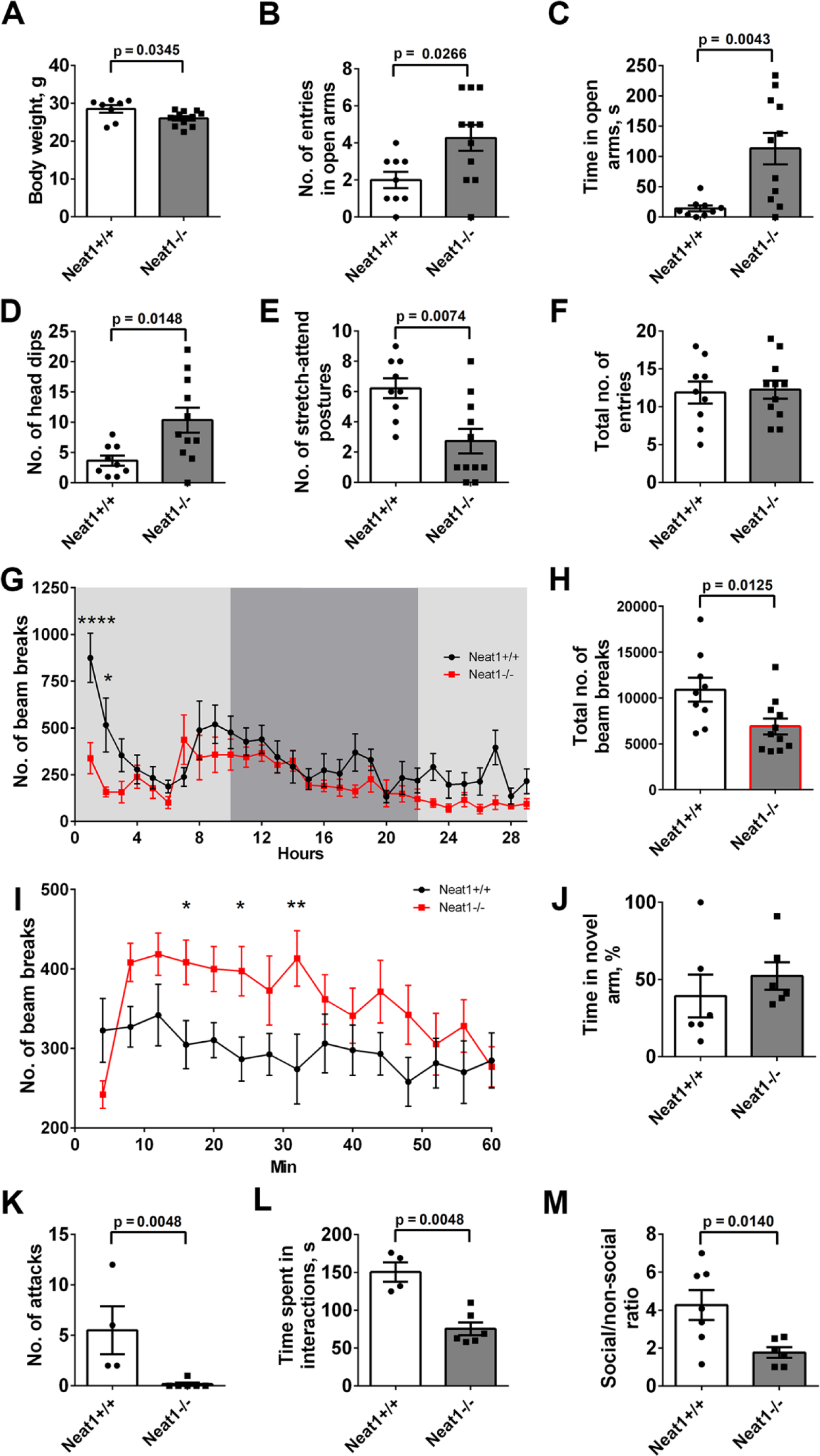
Behavioural alterations in *Neat1^-/-^* mice. (**A**) Decreased body weight in *Neat1^-/-^* mice (Mann-Whitney *U* test, effect size = 1.10; *Neat1^-/-^* n=8, *Neat1^+/+^* n=10). (**B-F**) Increased number of entries (**B**) and time spent in the open arms (**C**) as well as increased number of head dips (**D**) and stretch-attend postures (**E**) for *Neat1^-/-^* mice in elevated plus maze (EPM) (Mann-Whitney *U* test, effect size = 1.13, 1.54, 1.26, 1.42 respectively; *Neat1^-/-^* n=11, *Neat1^+/+^*n=9). Total number of arm entries was equal for the two genotypes (**F**). (**G, H**) Reduced locomotor activity of *Neat1^-/-^* mice in the Home Cage test (dim light conditions). Number of breaks per hour (**G**) and total number of breaks over the 30-h period (**H**) are shown (two-way ANOVA with Sidak’s multiple comparisons test in (**G**), ****p< 0.0001, *p< 0.05, main effect of group F (1, 522) = 38.90, effect size = 3.03; Mann-Whitney U test in (**H)**, effect size = 1.32; *Neat1^-/-^* n=10, *Neat1^+/+^*n=9). (**I**) Elevated locomotor activity of *Neat1^-/-^* mice under bright light conditions (activity chamber) (two-way ANOVA with Uncorrected Fisher’s LSD test, *p<0.05, **p< 0.01, main effect of group F (1, 261) = 26.58, effect size = 2.44; *Neat1^-/-^* n=11, *Neat1^+/+^* n=9). (**J**) Lack of working memory deficits in *Neat1^-/-^*mice. Intact short-term memory in *Neat1^-/-^* mice in the Y-maze test as indicated by the same percent of entries to novel arm (**K**), as compared to *Neat1^+/+^* mice (Mann-Whitney *U* test; *Neat1^-/-^* n=6, *Neat1^+/+^*n=6). (**K,L**) Decreased social interaction and aggression in *Neat1^-/-^* mice in the Resident-Intruder test as revealed by lack of attacks (**K**) and decreased time spent interacting with the intruder (**L**) (Mann-Whitney *U* test, effect size = 2.75 for both variable; *Neat1^-/-^* n=6, *Neat1^+/+^* n=4). (**M**) Decreased social interest in *Neat1^-/-^* mice revealed in the Social Odour test (Mann-Whitney *U* test, effect size = 1.82; *Neat1^-/-^*n=5, *Neat1^+/+^* n=6).

To analyse the locomotor activity of *Neat1^-/-^*mice under conditions of normal housing, we recorded their ambulatory activity for 24 h in the home cage (HC) test. *Neat1^-/-^* mice were overall hypoactive, with significantly reduced total activity during the test, as compared to *Neat^+/+^* mice (Fig. 2G,H). Decreased locomotor activity in the absence of motor impairment could be interpreted as depression-like behaviour. Surprisingly, *Neat1^-/-^*mice displayed dramatically reduced activity, as compared to *Neat^+/+^* mice, in the first two hours of the test, i.e. when the animals encountered novel conditions, suggestive of increased anxiety. We were initially puzzled by the discrepancies between locomotor activity and anxiety levels in *Neat1^-/-^* mice recorded in HC and EPM tests. However, there is one important difference between the two tests which is the brightness of light. HC test is conducted at 30-40 lux (during the day phase), which is comparable to the light intensity under normal housing conditions. In contrast, EPM test is carried out under the bright light conditions (∼400 lux). Thus in the EPM test, the mice were subjected to strong aversive stimulation – a combination of novelty and bright light, which could be the reason for abnormal behaviour of *Neat1^-/-^*mice. To test this directly, we measured locomotor activity under bright light conditions (600 lux) for 1 h in the activity cage apparatus. *Neat1^-/-^*mice were indeed hyperactive under these conditions (Fig. 2I). Therefore, in the absence of strong stress, *Neat1^-/-^*mice demonstrate decreased or similar (after habituation to novelty) level of locomotor activity to that of *Neat^+/+^* mice. However, a strong stressful stimulus, such as bright light, is sufficient to induce hyperactivity, impulsive reactions and possibly increased anxiety in *Neat1^-/-^* mice. Therefore, strong aversive stimulation converts the behavior of *Neat1^-/-^*mice from hypoactive to hyperactive. To rule out possible memory deficits which could affect EPM test performance, we examined working memory using Y-maze forced alternation task. This test did not reveal any differences between the genotypes (Fig. 2J and not shown).

To assess social behaviour of *Neat1^-/-^*mice, we used Resident-Intruder test. In this test, *Neat1^+/+^* mice demonstrated normal level of territorial defence behaviour, including moderately aggressive acts, such as lateral threats, chase and clinch attacks. In contrast, *Neat1^-/-^* mice showed nearly complete absence of aggression (Fig. 2K). In addition, *Neat1^-/-^*mice had decreased physical interactions with the intruder (Fig. 2L). Altered social behaviour was further confirmed in the Social Odour test, where *Neat1^-/-^* mice showed less interest in a social odour as compared to *Neat^+/+^* mice (Fig. 2M).

We next asked whether the phenotype in *Neat1^-/-^*mice would change during aging. Two key tests, EPM and HC, were performed in a group of 18-month old mice (novel cohorts of animals not subjected to behavioral testing before). In the EPM test, aged *Neat1^-/-^* mice showed behavioural patterns similar to the young animals, in that they also displayed preference for open arms and had increased number of head dips (fig S2A-D). Interestingly, *Neat1^-/-^* mice were also hyperactive in this test, as indicated by increased total number of entries (fig S2E). In the HC test, *Neat1^-/-^* mice demonstrated reduced activity in the first 4 h (statistical significance reached for the first hour), however their total activity was similar to that of *Neat^+/+^* mice (fig S2F,G). We also noticed an abnormal pattern of rhythmic fluctuations of activity in old *Neat1^-/-^* mice in this test: peaks of activity for *Neat1^-/-^* mice were often in the reverse phase to those of *Neat1^+/+^* animals (fig S2F). Young *Neat1^-/-^* mice also demonstrated smoothened rhythmic activity pattern after habituation (from 8 h onward), lacking bursts of activity alternating with quiet periods typical for *Neat^+/+^*mice (Fig. 2G).

To summarise, *Neat1^-/-^* mice display a distinct behavioural phenotype manifested in stress-induced hyperlocomotion, decreased anxiety, disinhibition, altered daily rhythms of activity and impaired sociability, without concomitant working memory or motor deficits, and this phenotype is largely preserved during aging.

### Neat1 knockout mice do not present with neuronal loss, neuroinflammation or synaptic pathology

We next sought to understand the mechanisms underlying the observed behavioural phenotype in *Neat1^-/-^* mice. We first asked if this phenotype could be of developmental origin. Using RNAScope® ISH, we detected almost zero Neat1 expression in the brain of E12.5 mouse embryos (fig S3). Similarly, in the original study, Neat1 was not detected in E9.5 mouse embryos (Nakagawa et al., 2011). Limited Neat1 expression during embryonic development of *Neat^+/+^* mice suggested that the observed behavioural phenotype in *Neat1^-/-^* mice is unlikely to be caused by developmental defects.

Analysis of the gross morphology of the brain of *Neat1^-/-^* and *Neat1^+/+^* mice using Nissl staining and NeuN immunohistochemistry did not reveal differences between the genotypes (Fig. 3A,B). Neuronal numbers in *Neat1^-/-^*mice were normal at least in the frontal cortex, consistent with the absence of overt neurological symptoms (Fig. 3C). However, we detected a small but significant decrease in the brain weight of *Neat1^-/-^*mice (Fig. 3D), which was in line with their reduced body weight (Fig. 2A).

**Figure 3.**
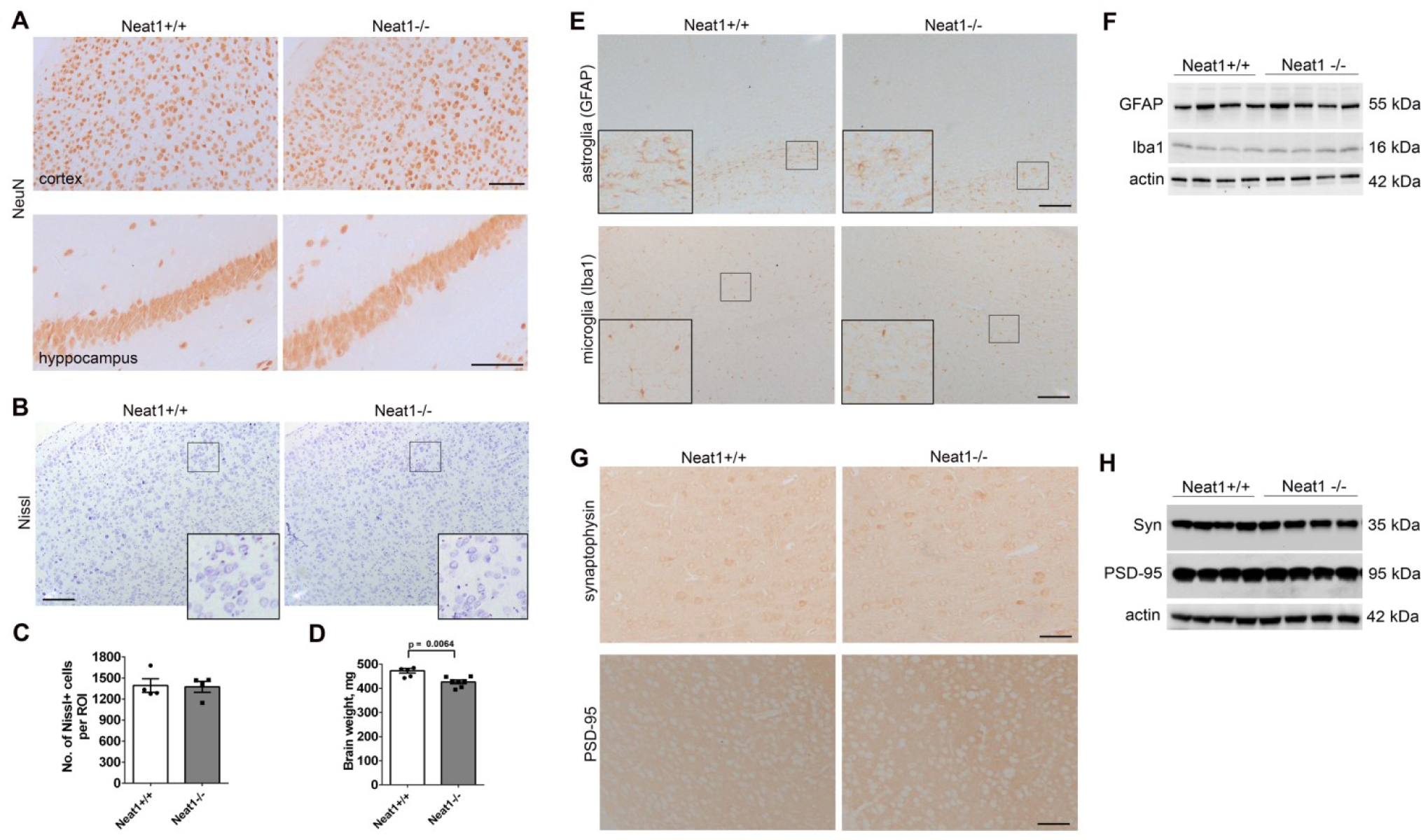
Loss of Neat1 does not induce neuronal loss, neuroinflammation or overt synaptic pathology. (**A**) Lack of gross changes in the anatomical structures of the brain in *Neat1^-/-^* mice as revealed by anti-NeuN staining. Representative images for the frontal cortex and hippocampus of 2-month old animals are shown. (**B,C**) Normal neuronal counts in the cortex of *Neat1^-/-^* mice. Representative images of Nissl-stained sections (**B**) and quantification (**C**) for 2-month old animals are shown (*Neat1^-/-^* n=4, *Neat1^+/+^* n=4). ROI, region of interest. (**D**) Reduced brain weight in *Neat1^-/-^* mice (Mann-Whitney *U* test, effect size = 2.15; *Neat1^-/-^* n=7, *Neat1^+/+^* n=6). (**E,F**) Lack of neuroinflammation in the brain of *Neat1^-/-^* mice. Distribution (**E**) and levels (**F**) of GFAP (astroglial marker) and Iba1 (microglial marker) were analysed in the cortex of *Neat1^-/-^* and *Neat1^+/+^*mice using immunohistochemistry and Western blotting, respectively. (**G,H**) Lack of obvious synaptic pathology in the brain of *Neat1^-/-^* mice. Distribution (**G**) and levels (**H**) of two main synaptic markers synaptophysin and PSD-95 were examined in the cortex of *Neat1^-/-^* and *Neat1^+/+^* mice using immunohistochemistry and Western blotting, respectively. In (**E**) and (**G**), representative images are shown. Scale bars, 100 µm.

We next examined levels of several markers commonly altered in models of neurological diseases. Neuroinflammation is typical for many neurological conditions, and Neat1 itself has been implicated in immune response (Imamura et al., 2014; Morchikh et al., 2017; Gast et al., 2019). However immunohistochemistry and Western blotting did not reveal any signs of astrogliosis or microgliosis in the brain of *Neat1^-/-^* mice (Fig. 3E,F). Synapse dysfunction is a major determinant of many neurological diseases. However distribution and levels of the two critical synaptic proteins, synaptophysin and PSD95 (encoded by *Dlg4* gene) in the brain of *Neat1^-/-^* mice appeared normal (Fig. 3G,H).

Therefore, behavioural deficits in *Neat1^-/-^*mice cannot be explained by neuronal loss, neuroinflammatory phenotype or gross synaptic dysfunction.

### Neat1 ^-/-^ neurons have altered calcium homeostasis and are hyperexcitable in vitro and in vivo

We speculated that abnormal behavioural responses of *Neat1^-/-^* mice may be determined by intrinsic changes in neurons, *e.g.* their altered excitability. It has been shown previously that human iPSC-derived neurons become hyperexcitable after acute downregulation of NEAT1 (Barry et al., 2017a). We prepared and analysed primary cultures of the hippocampal neurons from newborn *Neat1^+/+^* and *Neat1^-/-^*mice. Morphological analysis of neurons transiently transfected to express GFP did not reveal significant differences in the neurite outgrowth and branching between the genotypes (Fig. 4A). To assess excitability properties of these neurons, including transmembrane currents and the ability to generate induced action potentials (iAPs), we employed patch-clamp whole-cell electrophysiology. In current-clamp studies, 50% of *Neat1^-/-^*neurons demonstrated spontaneous action potentials, whereas in the *Neat1^+/+^*group, only 36% neurons were spontaneously active (Table S1). All examined *Neat1^-/-^*neurons displayed repetitive induced action potentials (iAPs “trains”) *vs.* only 40% of the neurons from the *Neat1^+/+^* group (Fig. 4B,C; Table S2). Another evidence of augmented activity of *Neat1^-/-^* neurons was that they demonstrated significantly higher frequency of train iAPs compared to *Neat1^+/+^*neurons (Fig. 4D). Interestingly, while having similar values of the resting membrane potential, *Neat1^-/-^* neurons had significantly faster depolarization and repolarisation rates (103.1 ± 18.4 V/s and −71.6 ±14.1 V/s *vs.* 61.6 ± 9.6 V/s and −37.9 ± 8.1 V/s for *Neat1^-/-^*and *Neat1^+/+^* neurons, respectively; p < 0.05).

**Figure 4.**
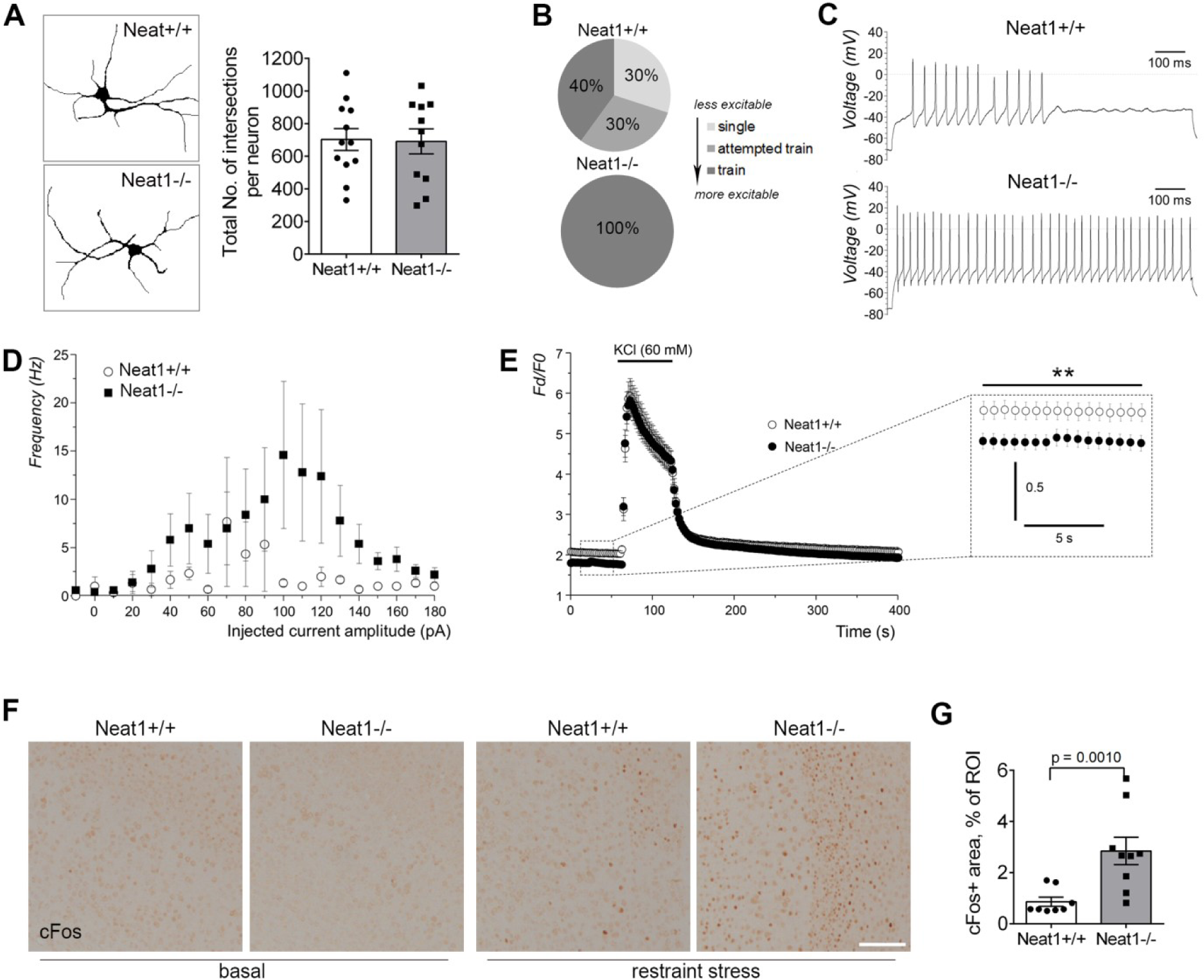
Neat1 knockout neurons are intrinsically hyperexcitable. (**A**) Primary hippocampal neurons isolated from *Neat1^-/-^*and *Neat1^+/+^* newborn mice do not display differences in cell morphology and neurite branching. Traces (left) and branching analysis (right) for DIV7 neurons are shown (*Neat1^-/-^* n=12, *Neat1^+/+^* n=12). Neurons were transiently transfected to express GFP and analysed 24 h post-transfection. (**B,C**) All *Neat1^-/-^* neurons generate trains of induced action potentials (iAPs). The proportion of neurons able to generate three patterns of induced activity (single action potential, attempted trains of action potentials or trains of action potentials) is shown (*Neat1^-/-^*n=4, *Neat1^+/+^* n=7) (**B**). Exemplar traces of current-clamp recordings when *Neat1^+/+^* (upper panel) and *Neat1^-/-^*(lower panel) neurons were held at between −70 and −80 mV and a following 1-s step of injected currents to the value that produced maximal trains of iAPs are also shown (**C**). (**D**) Representative spike frequency plots for *Neat1^+/+^* (open circles) and *Neat1^-/-^* (filled squares) neurons exhibiting trains of iAPs. (**E**) KCl-induced Ca^2+^ response in *Neat1^-/-^*and *Neat1^+/+^* neurons (two-way ANOVA with Fisher’s LSD test, **p< 0.01, main effect of group F (1, 1440) = 255.0, effect size = 4.82, **p<0.01; *Neat1^-/-^* n=27, *Neat1^+/+^* n=20). Inset shows the difference in the basal Ca^2+^ levels between the phenotypes. (**F,G**) Increased number of cFos-positive neurons in the cortex of *Neat1^-/-^* mice after the restraint stress (immobilisation). Representative images (**F**) and quantification (**G**) are shown (Mann-Whitney *U* test, effect size = 2.25; *Neat1^-/-^*n=3, *Neat1^+/+^* n=3; 9 ROIs). ROI, region of interest. Scale bar, 100 µm.

To understand the basis of the enhanced excitability of *Neat1^-/-^* neurons, transmembrane voltage-gated Na^+^ and K^+^ currents were measured using voltage-clamp mode. *Neat1^-/-^* neurons had increased densities for both inward and outward transmembrane currents with significant difference from *Neat1^+/+^*neurons for Na^+^ (p < 0.05) but not for K^+^ voltage-gated currents (fig S4A,B; Table S3). Activation/inactivation profiles of the voltage-activated Na^+^ currents for *Neat1^-/-^* neurons exhibited larger availability windows with high G/Gmax maxima and twice as much difference between mean half-activation and half-inhibition voltages (Va50 - Vi50) 5.0 ± 1.0 mV *vs*. the Neat1^+/+^ group 2.9 ± 1.1 mV. Vm values of both *Neat1^-/-^*and *Neat1^+/+^* neurons were falling within those windows however the proportion of spontaneously active neurons was higher in the *Neat1^-/-^*group (fig S4C,D; Table S1). The above data suggest that increased excitability of *Neat1^-/-^* neurons is determined by the increased transmembrane ion conductances particularly through Na^+^ voltage-gated channels, which are responsible for generation of neuronal action potentials.

To explore how the changes in excitability of the *Neat1^-/-^* neurons correspond to the regulation of calcium (Ca^2+^) homeostasis, we performed Ca^2+^ imaging experiments using ratiometric Fura-2 AM dye to examine the changes of the intracellular Ca^2+^ induced by KCl. Although Ca^2+^ responses upon KCl (60 mM) application in the neurons of both genotypes were similar, we found that *Neat1^-/-^*neurons have significantly lower (p<0.0001) basal levels of Ca^2+^ (Fig. 4E).

We next sought to confirm the increased activity of *Neat1^-/-^* neurons *in vivo*. The immediate early gene *cFos* is the most well-known molecular marker of neuronal activity (Zhang et al., 2002). Expression of this gene can be experimentally induced in the rodent brain *in vivo* by certain stresses such as immobilisation (restraint stress) (Melia et al., 1994). We subjected *Neat1^-/-^* and *Neat1^+/+^*mice to 60 min restraint stress followed by 60 min recovery and subsequently analysed cFos protein expression in the brain of these mice by immunohistochemistry. We found that the number of cFos-positive cells in the cortex of stressed *Neat1^-/-^* mice was significantly higher as compared to stressed *Neat^+/+^* mice (Fig. 4F,G).

Overall, our data indicate that *Neat1^-/-^*neurons are intrinsically hyperexcitable and have deficiencies in calcium homeostasis, which can result in abnormal gene expression responses *in vivo*.

### Loss of Neat1 perturbs local chromosomal environment and leads to subtle downregulation of multiple genes in the brain

Absence of detectable changes in the core CNS function-related markers (Fig. 3) suggested the existence of more subtle molecular changes that result in abnormal neuronal excitability and altered behavioural responses in *Neat1^-/-^* mice. In order to identify possible changes of the gene expression profiles in the CNS of *Neat1^-/-^* mice, we performed transcriptomic analysis of cortical tissue. Total RNA (RIN 8.7-9.2) from the whole cortices of 2-month old animals (four per genotype) was used. Prior to sequencing, Neat1 levels in these samples were measured by qRT-PCR. In the original study, total Neat1 (Neat1_1+Neat1_2) levels in mouse embryonic fibroblasts were found to be decreased to 6.0 ± 0.5%, whereas Neat1_2 was undetectable (Nakagawa et al., 2011). In the cortex of adult mice, total Neat1 was found to be decreased to 18.9 ± 0.02%, and Neat1_2 – to 8.2 ± 0.01% (Fig. 5A), probably reflecting tissue- and developmental stage-specific differences in the efficiency of *Neat1* silencing. Using Illumina NextSeq500 platform and paired-end sequencing, we obtained ∼50 million reads per sample. One of the *Neat^+/+^* samples showed abnormally low Neat1 expression and was identified as an outliner in the principal component analysis leading to its exclusion from the differential gene expression analysis. Examination of RNA-Seq peaks revealed that Neat1_1 is the main transcript expressed in the cortex, while Neat1_2 levels are very low in this tissue, consistent with previous reports (Nakagawa et al., 2011; Bluthgen et al., 2017). In accord with qRT-PCR data, RNA-Seq analysis detected significantly reduced yet still detectable levels of Neat1 in the cortex of *Neat1^-/-^*mice (Fig. 5B,C).

**Figure 5.**
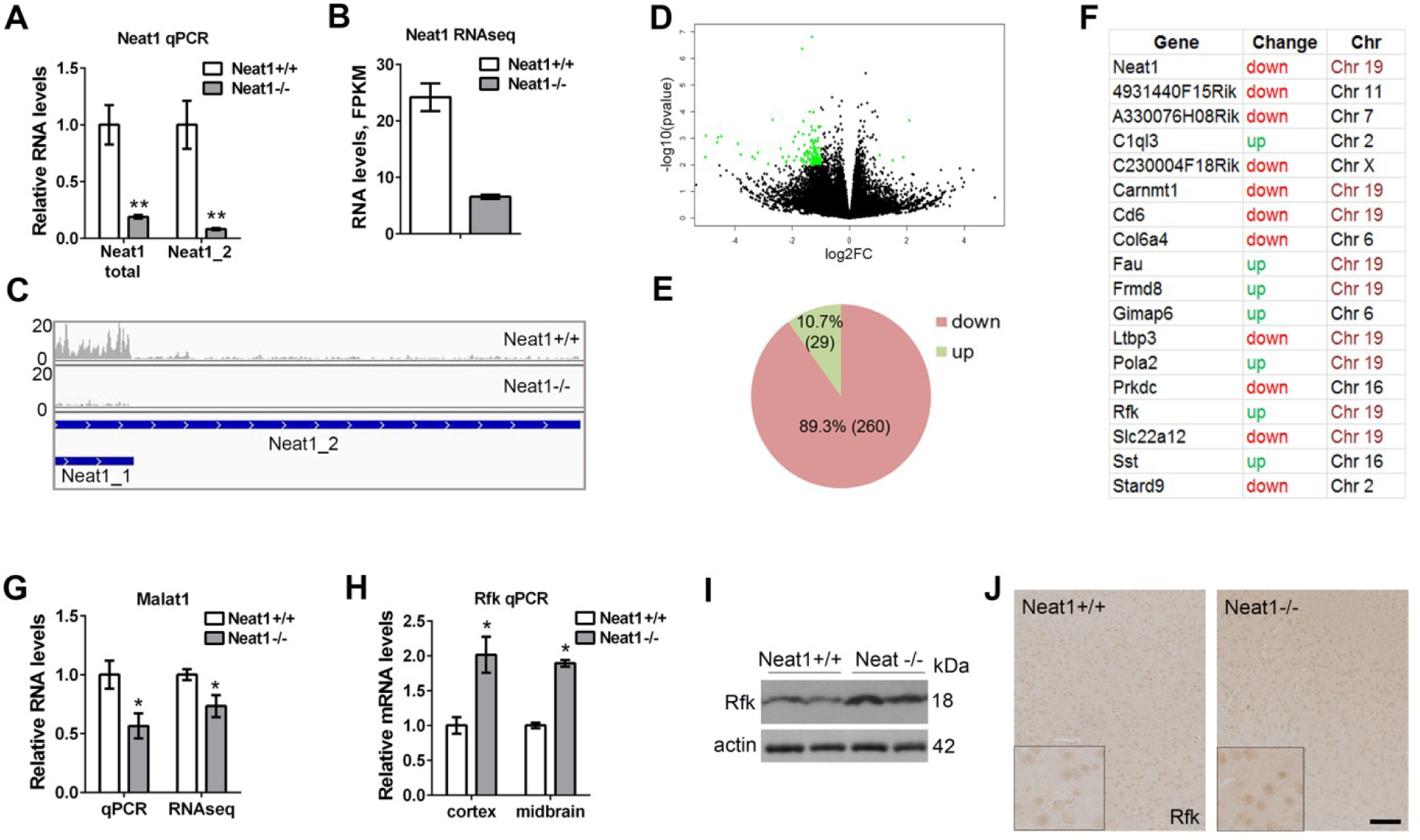
Loss of Neat1 results in subtle downregulation of multiple genes and significant expression changes of a small number of genes in the vicinity of the *Neat1* locus. (**A-C**) Significant downregulation of Neat1 isoforms in the cortex of *Neat1^-/-^* mice as revealed by qRT-PCR (**A**) and RNA-Seq (**B**, **C**). (Mann-Whitney *U* test, **p< 0.01; *Neat1^-/-^* n=4, *Neat1^+/+^* n=4). RNA-Seq peaks for the *Neat1* locus demonstrate preferential expression of Neat1_1 in *Neat1^+/+^* mice and residual Neat1_1 expression in *Neat1^-/-^*mice (**C**). (**D,E**) Subtle downregulation of multiple genes in the cortex of *Neat1^-/-^* mice. Volcano plot for *Neat1^-/-^ vs. Neat1^+/+^* mice is shown; genes with log2 fold change (log2FC)>1.5 and p-value<0.01 are highlighted in green (**D**). The proportion of up- and down-regulated genes is also shown (**E**, gene numbers are given in brackets). (**F**) List of annotated genes differentially expressed (padj<0.05) in both 1-month old and 2-month old *Neat1^-/-^*mice. Gene names were retrieved by mapping Ensembl gene IDs identified in DESeq2 analysis to the mouse genome using DAVID 6.8 database. Chromosomal location as well as direction of change are also given. (**G**) LncRNA Malat1 expressed from a locus adjacent to *Neat1* is downregulated in the cortex of *Neat1^-/-^* mice as revealed by qRT-PCR and RNA-Seq (Mann-Whitney *U* test, *p<0.05; Neat1^-/-^ n=4, Neat1^+/+^ n=4). (**H-J**) Upregulated expression of *Rfk* gene in the brain of *Neat1^-/-^*mice as demonstrated by qRT-PCR (**H**), Western blotting (**I**) and immunohistochemistry (**J**) (Mann-Whitney *U* test, *p<0.05; *Neat1^-/-^*n=4, *Neat1^+/+^* n=4). In (**J**), representative images of cortex are shown. Scale bar, 100 µm.

Differential gene expression analysis showed very subtle changes in the gene expression in *Neat1^-/-^* mice, consistent with the lack of overt neurological deficiencies in these mice. Only 14 genes were found to be significantly changed (adjusted p-value, padj<0.05), including ten genes with known function (*Cd6, Neat1, Rprm, Col62a, Prune2, Frmd8, Eps8l1, Crip2, Carnmt1, Snhg1*, Table S4). We next used the list of genes with at least 1.5-fold change in their expression (non-adjusted p-value<0.01), which was met by 289 genes, to assess gross gene expression changes. Overwhelmingly more genes from this list were found to be downregulated (260: 89.3%) than upregulated (29: 10.7%) (Fig. 5D,E), indicating that Neat1 positively regulates gene expression of a large subset of genes. Puzzled by this finding, we additionally performed a low coverage RNA-Seq analysis of another cohort of *Neat1^-/-^* and *Neat1^+/+^*animals (aged 1 month). Similar to the 2-month old mice, changes in the gene expression were very subtle, however, the vast majority (85%) of differentially expressed genes (DEGs: >1.5 fold change, p-value<0.05) were also found to be downregulated. In this latter mouse cohort, only four genes showed significantly altered expression (padj <0.05), namely *Neat1*, *Frmd8*, *Carnmt1* and *Malat1*. We compared the lists of genes whose expression was changed >1.5 fold in the cohorts of 1-month old mice (cut-off p-value <0.05, n=474) and 2-month old mice (cut-off p-value <0.01, n=289). Altogether, 29 genes, 18 of which are genes with known function, were found to appear in both groups, including previously identified genes with robust changes - *Neat1*, *Frmd8* and *Cd6* (Fig. 5F). Positional analysis for these genes showed that half of them (9/18) are located on chromosome 19, and some of them – in close vicinity to the *Neat1* locus (*Frmd8, Ltbp1, Pola2, Fau),* likely reflecting changes in the chromosomal environment upon *Neat1* disruption (Fig. 5F). However, expression of other genes surrounding the *Neat1* locus (*Scyl1*, *Slc25a45*, *Tigd3*, *Dpf2,* and *Cdc42ep2*) was not altered.

We next performed enrichment analysis on the set of genes with at least 1.5 fold change in their expression (p-value <0.01) for 2-month old mice. Out of these 289 mouse Ensembl gene IDs, 60 could be mapped to known genes using DAVID 6.8 tool. Small enrichment in anion transport genes (GO:0006820, p=0.00678: *Gabrr2, Anxa1, Kcnj10, Slc2a2* and *Slc22a12*) was identified, of which one gene, *Gabrr2*, attracted our attention. Stress-induced impulsive behaviours in animals may be suggestive of abnormal inhibitory neurotransmission, which is mediated by GABA receptors. We therefore examined the expression of *Gabrr2* and two major GABA receptor subunit genes, *Gabra1* and *Gabra2* by qRT-PCR. Although *Gabrr2* mRNA was indeed confirmed to be downregulated, *Gabra1* and *Gabra2* expression was unaltered (fig S5A). Further, numbers and distribution of inhibitory Gad67-positive interneurons in the brain as well as levels of Gad67 in the cortex were not changed in *Neat1^-/-^* mice (fig. S5B,C). Thus we concluded that GABAergic inhibitory transmission is unlikely to be significantly affected in *Neat1^-/-^* mice.

One gene found to be significantly changed (padj<0.05) in 1-month *Neat1^-/-^* mice was *Malat1*. Malat1 is a conserved lncRNA transcribed from a locus adjacent to *Neat1* and highly expressed in the brain (Zhang et al., 2012). *Malat1* null mice have also been reported to have upregulated expression of several genes in close vicinity to the *Malat1* locus, including *Neat1* and *Frmd8*, in the liver and brain (Zhang et al., 2012). However, another *Malat1* knockout mouse strain displayed decreased Neat1 expression in a subset of tissues, although brain was not examined (Nakagawa et al., 2012). Our RNA-Seq analysis showed Malat1 downregulation in *Neat1^-/-^* mice, in both age groups analysed, and this result was confirmed by qRT-PCR (Fig. 5G).

Another gene found significantly upregulated in both 1- and 2-month old *Neat1^-/-^* mice encodes riboflavin kinase (Rfk) enzyme. Intriguingly, human riboflavin kinase has been recently linked to neurological diseases (Johnson et al., 2012) and regulation of circadian rhythms (Hirano et al., 2017). We confirmed that Rfk mRNA is ∼2 fold upregulated in the cortex and midbrain of *Neat1^-/-^* mice by qRT-PCR (Fig. 5H). Consistent with this, Rfk protein levels were higher in the cortex of *Neat1^-/-^* mice, as analysed by Western blot (Fig. 5I). Immunohistochemical analysis revealed exclusively neuronal Rfk expression, and Rfk upregulation was also evident in the brain sections of *Neat1^-/-^* mice (Fig. 5J).

In sum, Neat1 acts as a weak positive modulator of gene expression in the brain, whilst its loss impacts on the transcription of a small pool of genes in the vicinity of the *Neat1* locus.

### Loss of Neat1 affects alternative splicing of genes involved in neuronal homeostasis

RNA-Seq datasets for *Neat1^-/-^* and *Neat1* ^+/+^ mice were further used to analyse possible changes in the alternative splicing in the brain of *Neat1^-/-^* mice. For that, a recently developed pipeline, LeafCutter (Li et al., 2018b), was utilised. Altogether, 14 differentially spliced genes were identified in *Neat1^-/-^*mice using a < 20% FDR threshold (Fig. 6A,B and Table S5). All these genes are highly expressed in the brain and two of them – almost exclusively in the brain (*Myt1l, Rgs7*). Most of these genes are also important for the CNS function, including those with roles in neuron proliferation, differentiation, cell-cell interaction, intraneuronal transport, synaptogenesis, primary axonogenesis, and axon guidance (*Dlg1, Myt1l, Ptprs, Dctn2, Rgs7*). Further, some genes are genetically linked to psychiatric disorders such as schizophrenia (*Dlg1, Myt1l*) (Lee et al., 2012; Uezato et al., 2017), autism spectrum disorder (*Myt1l*) (Meyer et al., 2012; Wang et al., 2016) and panic disorder (*Rgs7*) (Hohoff et al., 2009). Other genes from this list are linked to various types of cancer (*Pnck, Calu, Vav2, Kif3a, Adgrf5, N4bp2l2*).

**Figure 6.**
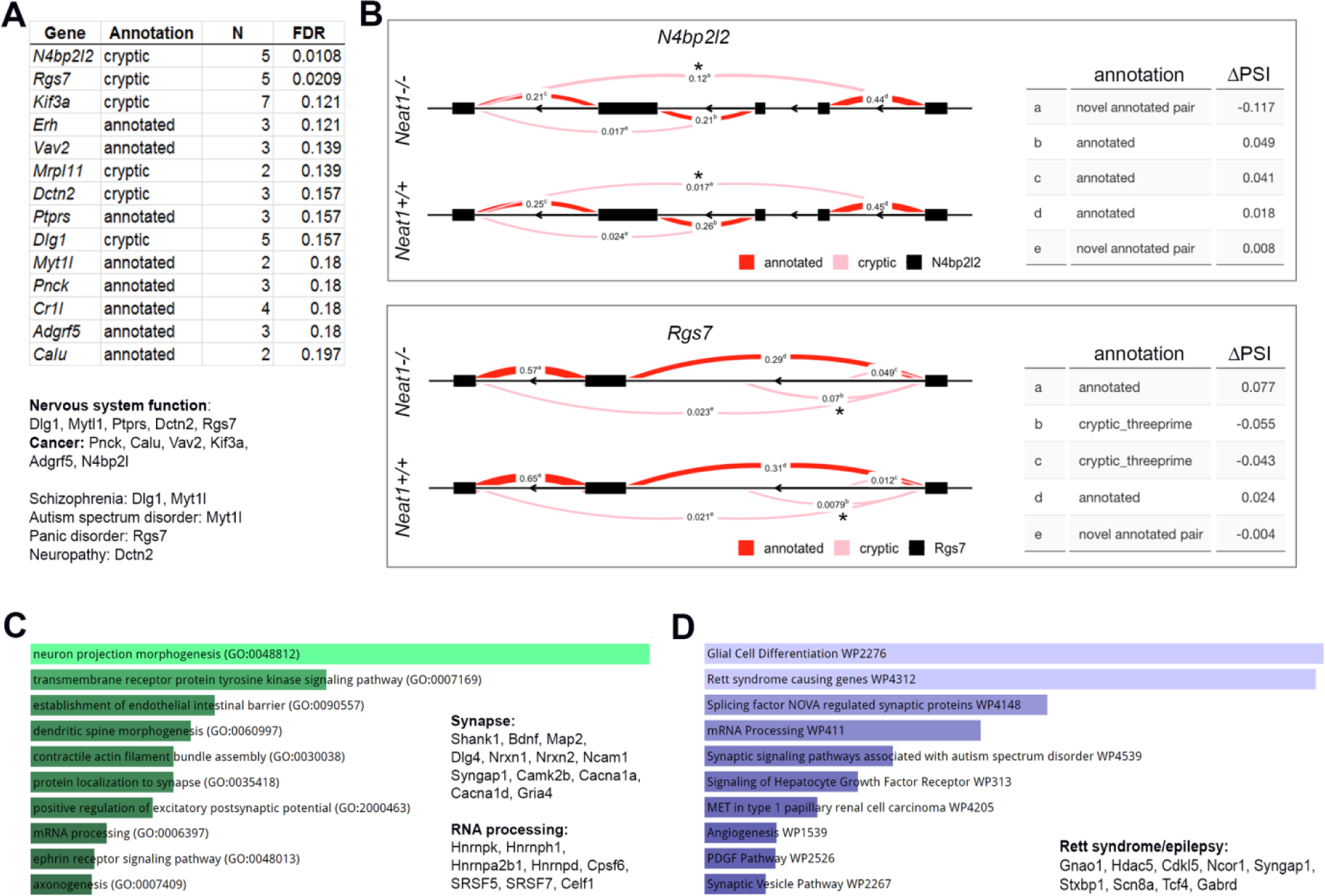
Loss of Neat1 causes changes in the alternative splicing of genes involved in synaptic function and RNA processing. (**A**) List of genes differentially spliced in the cortex of *Neat1^-/-^* mice. Functional association and role in disease for these genes are also given. N, number of introns in a cluster; FDR, false discovery rate. (**B**) Splicing diagrams showing differential intronic usage in two intron clusters of *N4bp2l2* and *Rgs7* genes in *Neat1^+/+^* and *Neat1^-/-^*mice. Intron usage is indicated by Percent Spliced In (PSI). Table on the right lists changes in PSI (ΔPSI). (**C,D**) Enrichment analysis of differentially spliced genes as analysed using Enrichr. GO term Biological Process (**C**) and Wiki Pathways (**D**), alongside with their functional association and role in disease are shown.

For the enrichment/pathway analysis we used a p-value cut-off of <0.05 which was met by 671 genes (Table S5). The use of a relaxed cut-off (p-value instead of padj) in this case was possible because only truly positives are expected to cluster in functional categories whereas false positives should be functionally unrelated (Cooper-Knock et al., 2012). Analysis of GO Biological process enriched terms using Enrichr revealed that a significant number of differentially spliced genes were involved in axonogenesis, protein localization to synapse and dendritic morphogenesis, including genes critical for neuronal homeostasis and implicated in neurological diseases, e.g. *Shank1, Bdnf, Map2, Dlg4, Nrxn1, Nrxn2* and *Ncam1* (Fig. 6C). Another enriched category was RNA processing, including genes encoding abundant RNA-binding proteins *Cpsf6, Hnrnpk, Hnrnph1, Hnrnpa2b1, Hnrnpd, Srsf5* and *Srsf7*. Pathway analysis of this dataset also identified genes involved in synaptic function (e.g. *Syngap1, Camk2b, Cacna1a, Cacna1d, Gria4*) and mRNA processing (e.g. *Celf1, Cstf2, Pcbp2, Hnrnpk, Hnrnph1, Hnrnpa2b1*, alongside with several genes encoding serine and arginine splicing factors) (Fig. 6D). Multiple genes from this list are linked to neurological conditions. Calcium channel subunits *Cacna1d* and *Cacna1a* are associated with autism spectrum disorders (Limpitikul et al., 2016) and with familial hemiplegic migraine and episodic ataxia (Grieco et al., 2018; Jen and Wan, 2018), respectively. Variants in the *Gria4* gene are linked to schizophrenia and intellectual disability (Makino et al., 2003; MacDonald et al., 2015; Martin et al., 2017). Consistent with recently reported dysregulation of NEAT1 in epilepsy (Lipovich et al., 2012; Barry et al., 2017a; Bluthgen et al., 2017), the second enriched pathway in our analysis was “*Rett syndrome causing genes*” which contained 9 genes implicated in epilepsy, such as *Gabrd, Gnao1, Stxbp1* and *Scn8a*.

Overall, we showed that loss of Neat1 perturbs alternative splicing in the brain, with the most significant effect on the genes related to synaptic function and RNA metabolism and implicated in neurological diseases.

## Discussion

LncRNAs play an important role in fine-tuning the functions specific for different cell types, and their activity might be especially important in the adult nervous system. Indeed, lncRNAs have been reported to regulate critical processes in the developing and adult brain, including cell differentiation, neurite elaboration and synaptogenesis (Briggs et al., 2015; Wang et al., 2017). NEAT1 is an abundant lncRNA known to be expressed in the mammalian CNS however its role in the CNS under basal conditions and under conditions of physiological stress remained largely unknown.

In the current study we performed a detailed behavioural characterisation of the Neat1 knockout mouse line which revealed a distinct phenotype whose core feature is the exaggerated response to physiological stress. Under stressful conditions, these mice adopt an active coping strategy (“fight-or-flight” response), instead of passive coping strategy or freezing typical for WT mice. In addition, *Neat1^-/-^* mice were found to have deficiencies in social interaction – encounter with unfamiliar animal or its odour, i.e. acute social stress. Furthermore, these mice also present with abnormal rhythmic fluctuations of daily activity, which is in line with the recently reported role for Neat1 in the control of circadian genes (Torres et al., 2016). Behavioural features detected in these mice, i.e. hyperactivity, reckless behaviours, disinhibition, impulsivity, are commonly found in psychiatric disorders, including dementia, bipolar disorder, schizophrenia, and attention deficit hyperactivity disorder (ADHD) (Russell, 2011; Yen et al., 2013; Beyer and Freund, 2017). Consistently, altered NEAT1 expression is a common theme in multiple neurological conditions including dementia (Tollervey et al., 2011), Alzheimer’s disease (Puthiyedth et al., 2016), Huntington’s disease (Sunwoo et al., 2016), schizophrenia (Li et al., 2018a; Katsel et al., 2019) and epilepsy (Lipovich et al., 2012; Barry et al., 2017a). Therefore, abnormal NEAT1 expression and/or regulation might underlie at least some of the symptoms and phenotypes in common neurological diseases.

Our study is the first to report the analysis of Neat1 expression and distribution in the mammalian CNS regions using *in situ* hybridization. We found that in the murine CNS (adult brain and spinal cord), Neat1 is expressed in multiple cell types with higher levels in glial cells. An earlier transcriptomic analysis of purified cell types also reported higher Neat1 levels in glial cells – astrocytes and microglial cells (Dong et al., 2015). In another study, NEAT1 was also reported to be highly expressed in oligodendrocytes (Katsel et al., 2019).

The longer isoform of Neat1, Neat1_2, is the main structural component of paraspeckles (Sunwoo et al., 2009). However, Neat1_2 expression is restricted to certain organs/tissues, such as the gut, and was not detected in the CNS (Nakagawa et al., 2011). Using RNA-Seq, we also show that Neat1_2 expression is barely detectable in the brain under basal conditions. Therefore, the phenotypes of *Neat1^-/-^* mice detected in the current study can be largely ascribed to the loss of Neat1_1 expression. However, a previous study demonstrated transient accumulation of Neat1_2 in the murine hippocampus after seizures (Bluthgen et al., 2017), suggesting that the isoform switch can take place in neurons under certain conditions. NEAT1_2/paraspeckles have a prosurvival role in several contexts and might contribute to neuroprotection in neurodegenerative diseases such as ALS (An et al., 2018). It will be important to establish whether bursts of Neat1_2 production may occur in the CNS and, more specifically, in neurons *in vivo* in response to various physiological and pathological stresses.

Although NEAT1 has been heavily implicated in the function of the immune system and in inflammatory response (Imamura et al., 2014; Morchikh et al., 2017; Gast et al., 2019), we show that its loss in mice does not result in a neuroinflammatory phenotype. This result is consistent with the role for NEAT1 as a stress-responsive transcript which would mediate/facilitate a response to an external trigger (pathogen) but likely does not have housekeeping immune functions.

It has long been known that deficits in the inhibitory GABAergic neurotransmission lead to behavioural changes in mice characterized by hyperactivity, panic attacks, impulsivity and generalization of fear. Therefore, it was possible that the hyperactive phenotype detected in *Neat1^-/-^*mice is caused by attenuated GABA system function. However, in RNA-Seq analysis, we found that only one member of GABA type A receptor complex *Gabrr2* is downregulated and another one, *Gabrd,* is differently spliced in the cortex of *Neat1^-/-^* mice. Immunohistochemistry and Western blot analysis also did not reveal any detectable changes in the populations of GABAergic neurons throughout the brain of *Neat1^-/-^* mice. While it cannot be excluded that subtle changes in the GABAergic neurotransmission are still present in *Neat1^-/-^* mice, we concluded that it is not the main driver of the observed behaviour changes. Another possibility was that neurons derived from *Neat1^-/-^* mice are intrinsically hyperexcitable, which was indeed the case and in line with previous data obtained in human neurons with acutely downregulated NEAT1 (Barry et al., 2017a). Augmented neuronal activity upon depletion of NEAT1 was previously determined indirectly, using calcium measurements. In our study, we used the gold standard method for this type of studies – whole-cell patch clamp recording in cultured neurons – which allowed to unambiguously establish the enhanced activity of *Neat1^-/-^*neurons. Comparison of the electric properties showed that increased excitability of *Neat1^-/-^* neurons was likely due to the upregulated voltage-gated Na^+^ influx, which determines the kinetics of the action potential and frequency of firing. Two-fold enlargement of voltage range availability for Na^+^ current in *Neat1^-/-^*neurons presumably allows greater number of these neurons to generate spontaneous and induced action potentials, whereas a trend for enhanced K^+^ conductance *Neat1^-/-^* neurons could be crucial for faster repolarization and therefore shortening of the absolute refractory period. Finally, we were able to demonstrate the relevant consequence of enhanced neuronal excitability – increased stress-induced cFos expression – in live mice. Our *in vivo* cFos data are consistent with a recent finding that Neat1 inhibits the expression of *cFos* in cultured neurons (Butler et al., 2019). Notably, basal cFos expression was not changed in the brain of *Neat1^-/-^*mice in our RNA-Seq analysis, therefore cFos regulation by Neat1 mainly occurs upon stimulation, at least *in vivo*.

In an attempt to delineate molecular (gene expression) changes which underlie the hyperexcitability phenotype in *Neat1^-/-^*neurons, we analysed differential gene expression and changes in the alternative splicing in the cortex of *Neat1^-/-^* mice. DEG analysis revealed only a small number of significantly changed genes, and prominent changes were found in the expression of genes located in the vicinity of *Neat1* such as *Frmnd8*, *Cd6* and *Malat1,* similar to a previous study (Katsel et al., 2019). This suggests that lncRNAs like NEAT1 and MALAT1 are important for regulation of gene expression in *cis*. Unexpectedly, we found that the majority of DEGs (∼90%) in *Neat1^-/-^* mice were downregulated, and this finding was further reproduced in an independent mouse cohort. Both NEAT1 and MALAT1 have been reported to be localised to numerous genomic sites and bind active genes (West et al., 2014). Thus Neat1 likely contributes to positive regulation of gene expression, explaining why its loss results in attenuated transcription of multiple targets. It is conceivable that moderate Malat1 downregulation found in *Neat1^-/-^* mice synergises with Neat1 loss to mediate this phenotype. In future studies, it will be important to characterise possible neurological changes in Malat1 knockout mice at the behavioral and molecular levels.

Alternative splicing occurs at high frequency in the CNS and is crucial for normal functioning of the brain (Yeo et al., 2004). Indeed, global changes in alternative splicing pattern have been identified in the brain of patients with psychiatric diseases such as schizophrenia (Reble et al., 2018). In our RNA-Seq dataset, we also detected a number of genes differentially spliced in the cortex of *Neat1^-/-^*mice. Functionally, these genes can be subdivided into groups related to *synaptic functions, mRNA processing* and *cancer*. Even though we did not detect major synaptic abnormalities in *Neat1^-/-^* mice, it is plausible that these alternative splicing changes translate into functional changes in the synapse and contribute to the observed behavioural phenotype. The role of lncRNAs in neural plasticity is emerging, the latter being crucial for stress-coping mechanisms and behavioural adaptation. For example, in response to repeated social defeat stress, hundreds of lncRNAs change their expression in the prefrontal cortex in mice, and these lncRNAs are involved in the regulation of synaptic function (Wang et al., 2019). NEAT1 interacts with multiple RNA-binding proteins (Nakagawa et al., 2018) many of which are autoregulated, including via alternative splicing (Jangi et al., 2014). Therefore, Neat1 loss can also lead to dysregulation of large RNA-binding protein networks. Finally, the pathophysiological connection between Neat1 isoform expression and tumorigenesis is well-established, and our data suggest that at least some of the cancer-related network changes caused by altered Neat1 expression may be attributed to abnormal alternative splicing regulation (Liu and Cheng, 2013). Further studies are needed to determine whether the effects of Neat1 loss on gene expression and alternative splicing are specific for the CNS tissue.

There are several limitations to our study. Firstly, the expression of Neat1 is not completely abolished in the mouse line studied, therefore it allowed us to characterise the consequences of its substantial downregulation, rather than its complete loss. Secondly, we cannot exclude possible effects of developmental compensation of Neat1 function due to the constitutive type of the knockout model. Finally, in our studies, we have not addressed the contribution of the two Neat1 transcripts to the observed phenotype. Despite very low Neat1_2 expression in the brain, it is still possible that its expression is transiently activated under stress and at least partially mediates stress-induced signaling in neurons. Thus, in future studies, it would be important to recapitulate our findings in tissue-specific conditional knockout models of Neat1 loss of function, including those with differential expression of the two isoforms.

Unlike CNS-specific lncRNAs, Neat1 is expressed in a variety of tissues. Presumably, such lncRNAs play a universal regulatory role, while the specificity arises from the repertoire of cell type-depended targets and environmental cues. It can be concluded that although Neat1 is not essential for basic brain functions, its availability becomes important in the presence of biologically significant stress stimuli. Overall, Neat1 plays a fine-tuning regulatory role in the nervous system and participates in the processes responsible for environmentally appropriate behaviour.

## Acknowledgments

The study was supported by fellowships from Medical Research Foundation and Motor Neurone Disease Association (Shelkovnikova/Oct17/968-799) to TAS. HA is a recipient of Cardiff University/China Scholarship Council PhD studentship. We thank Polina Yarova for the help with calcium imaging. DNA sequencing was undertaken at the Cardiff University School of Biosciences Genomics Research Hub.

## Supplementary Materials

**Fig. S1.**
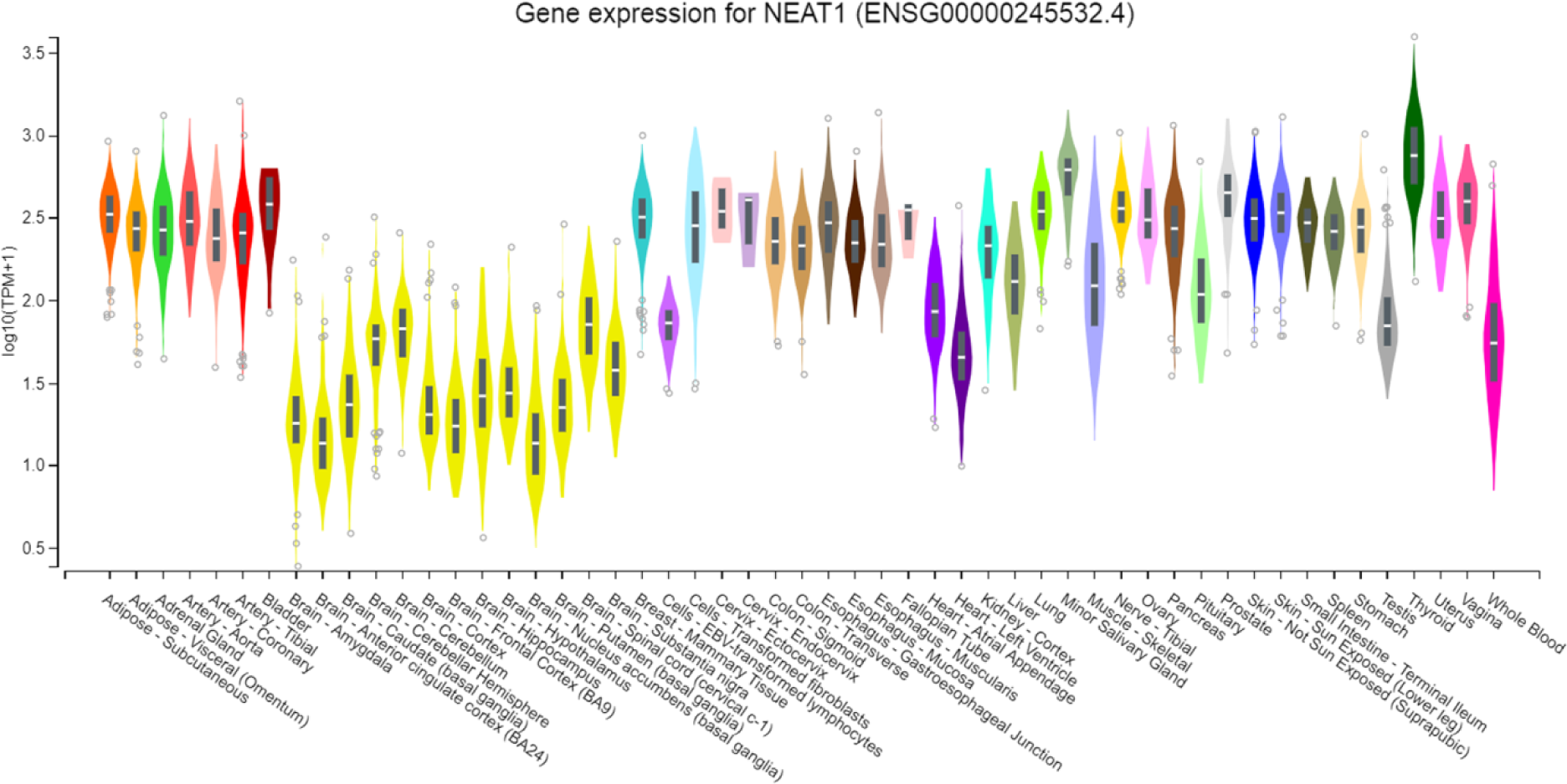
NEAT1 expression in the human organs and tissues according to the Genotype-Tissue Expression (GTEx) database (https://gtexportal.org/home/).

**Fig. S2.**
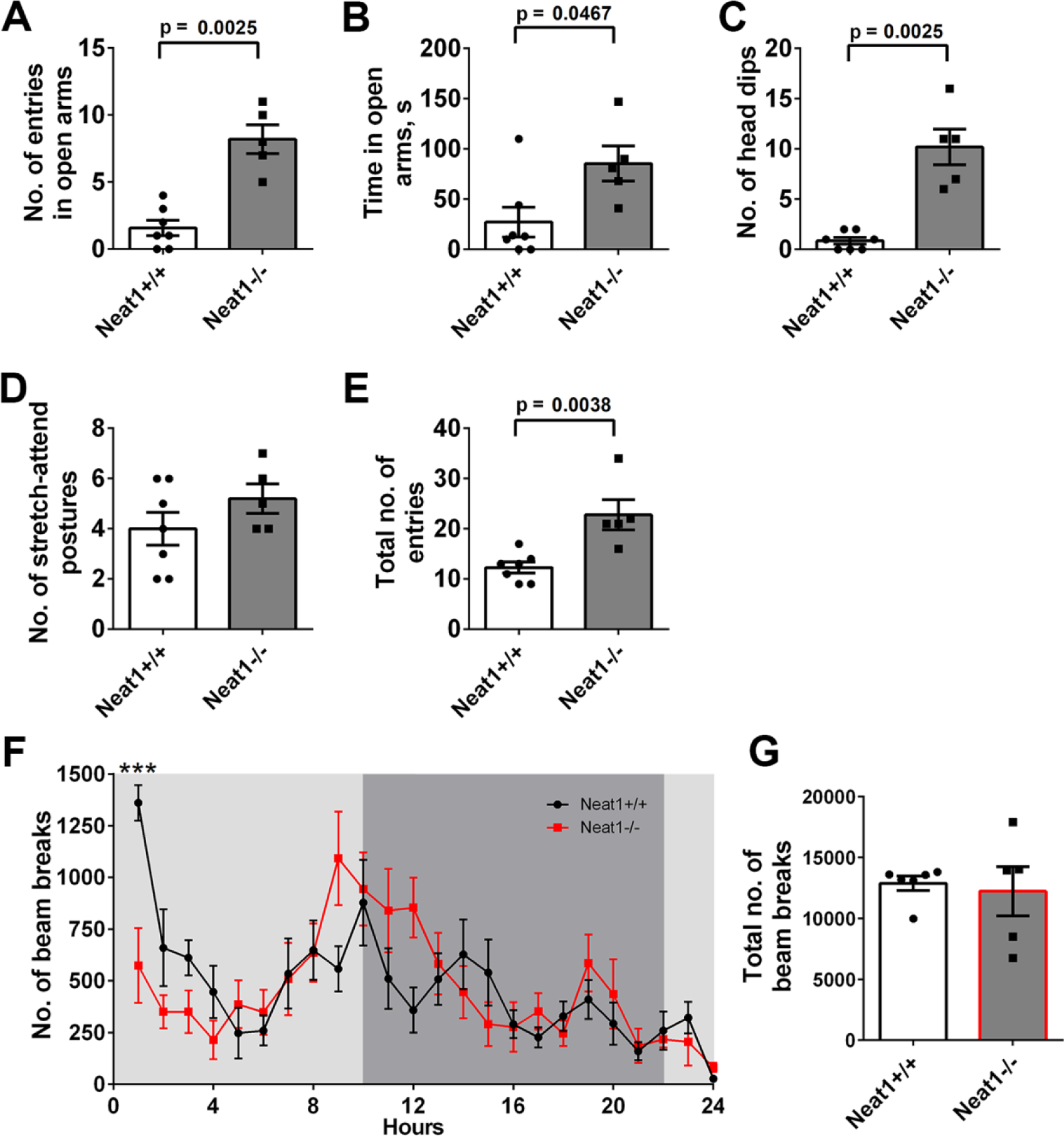
The behavioural phenotype of *Neat1^-/-^* mice is preserved during aging. Novel cohorts of 18 month-old animals not previously subjected to behavioural testing were used in these tests. (**A-E**) Performance of *Neat1^-/-^* and *Neat1^+/+^*mice in the elevated plus maze (EPM) test. Increased number of entries (**A**), increased amount of time spent in the open arms (**B**), increased number of head dips (**C**), unaltered number of stretch-attend postures (**D**) and increased total number of entries (**E**) for aged *Neat1^-/-^* mice as compared to *Neat1^+/+^*mice (Mann-Whitney *U* test, effect size = 2.87, 1.45, 2.87, 2.44 respectively; *Neat1^-/-^* n=5, *Neat1^+/+^* n=7). (**F,G**) Reduced locomotor activity of *Neat1^-/-^* mice in the Home Cage test during habituation (two-way ANOVA with Sidak’s multiple comparisons test, ***p< 0.001, main effect of group F (1, 216) = 0.004815, effect size = 2.93; *Neat1^-/-^*n=5, *Neat1^+/+^* n=6). Number of breaks per hour (**F**) and total number of breaks over the 24-h period (**G**) are shown.

**Fig. S3.**
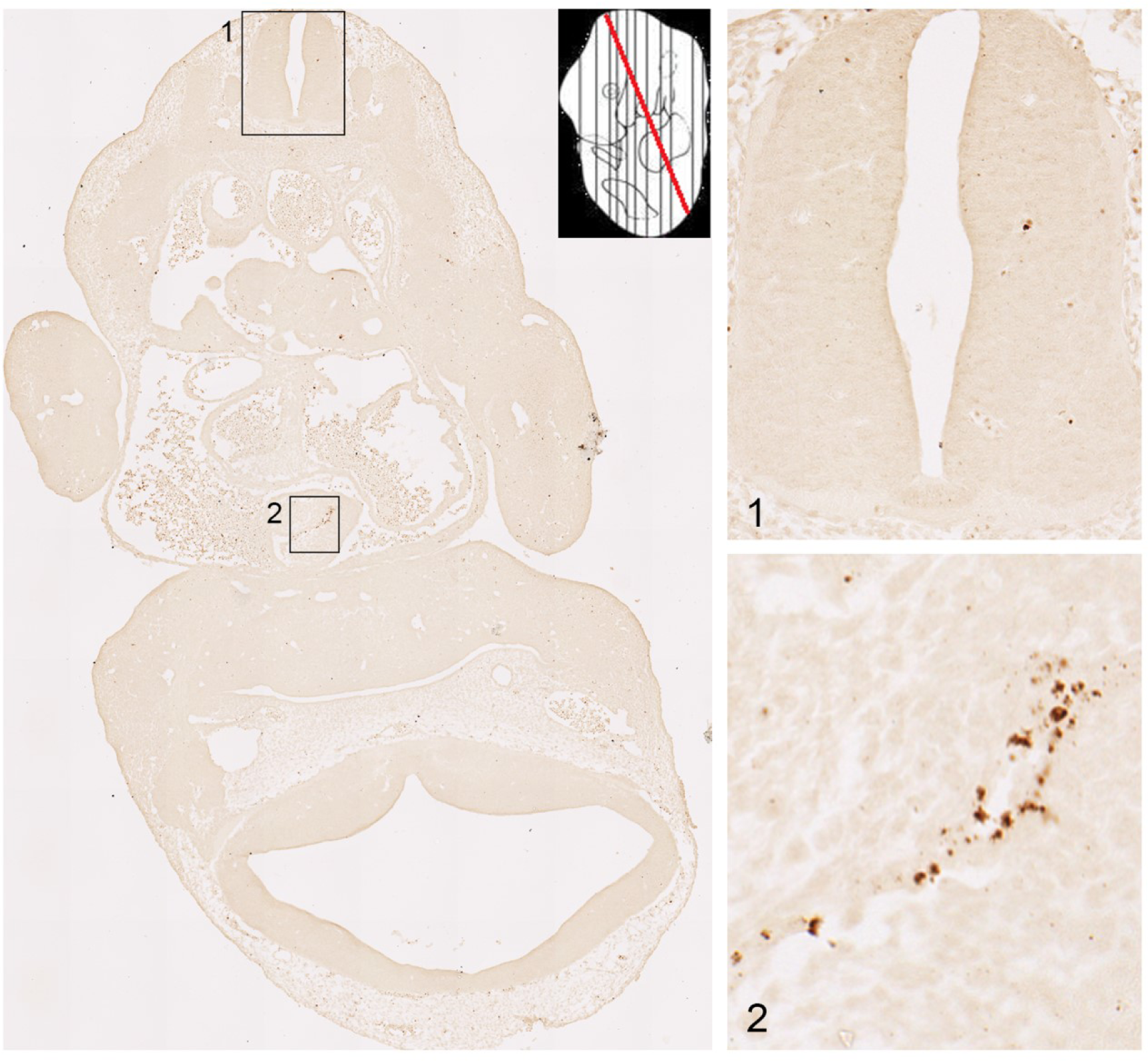
Neat1 expression in the E12.5 mouse embryo. RNAScope ISH using 5’ fragment Neat1 probe in WT E12.5 mouse embryo. The plane of the section within the embryo is also shown (red line). Numbered insets show two zones at higher magnification: 1 – neural tube, 2 – blood vessel. Note multiple Neat1-positive cells in the blood vessel but not in the neural tube.

**Fig. S4.**
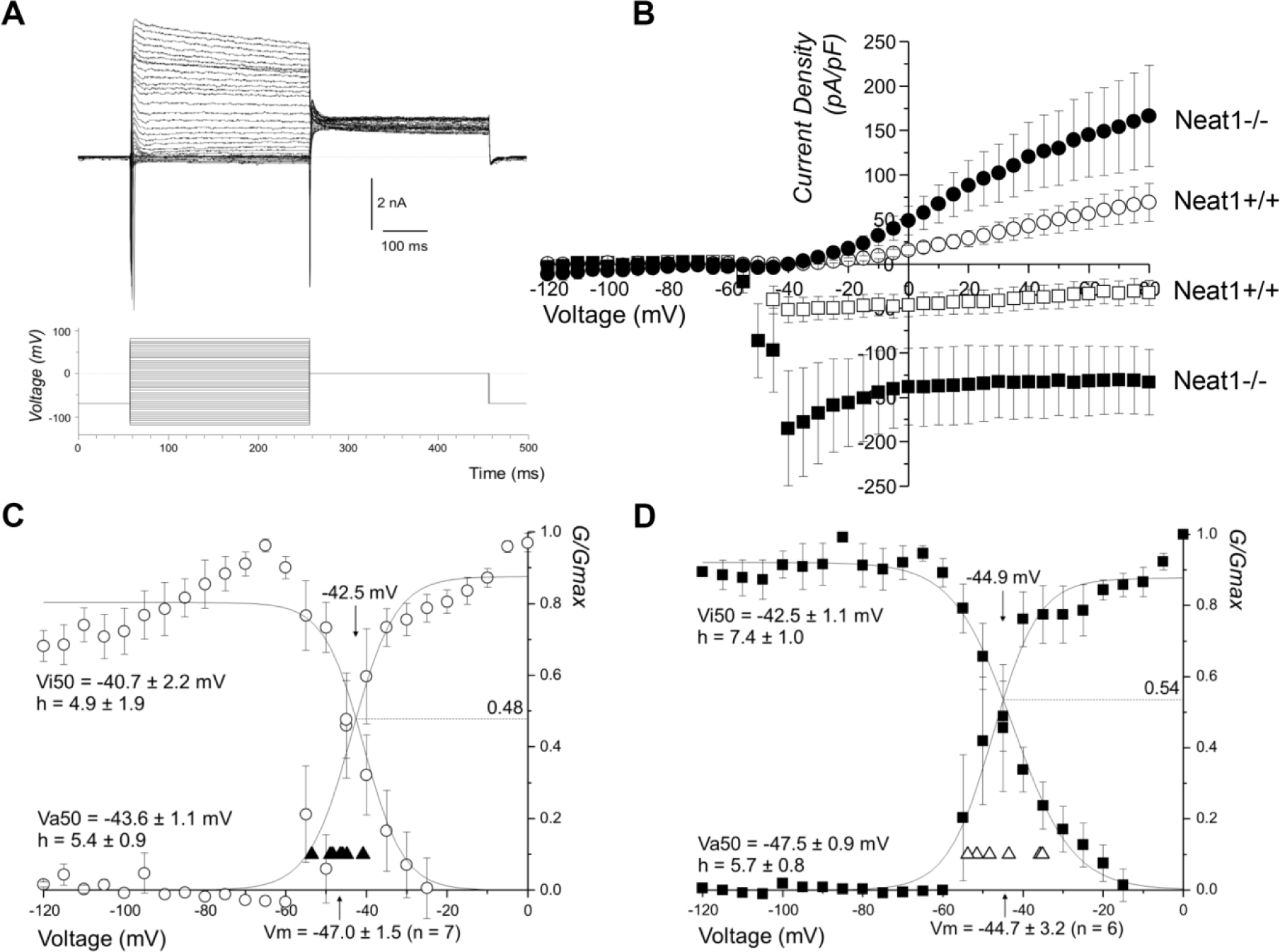
The effect of Neat1 loss on transmembrane voltage-gated K+ and Na+ currents in mouse neurons. (**A**) Exemplar families of whole cell currents evoked by the voltage activation/inactivation protocol (as shown at the bottom panel) in mouse neurons. (**B**) Mean current densities vs. voltage plots derived for voltage-activated Na+ (squares) and K+ currents (circles) derived from traces exemplified in (**A**). Empty and filled symbols represent the current densities from *Neat1^+/+^* and *Neat1^-/-^* mice, respectively. (**C-D**) Mean fractional conductance (G/Gmax) plots for voltage activation and inactivation of Na^+^ currents for *Neat1^+/+^* (**C**) and *Neat1^-/-^* (**D**) neurons derived from the traces exemplified in (**A**). On each panel are also shown individual Vm values (triangles) and mean Vm values (arrow on abscissa). Voltages of half-maximal action (Va50) and half-maximal inactivation (Vi50) are also indicated, along with h factors, mean crossing points (downward arrows) number of cells recorded for each group (n).

**Fig. S5.**
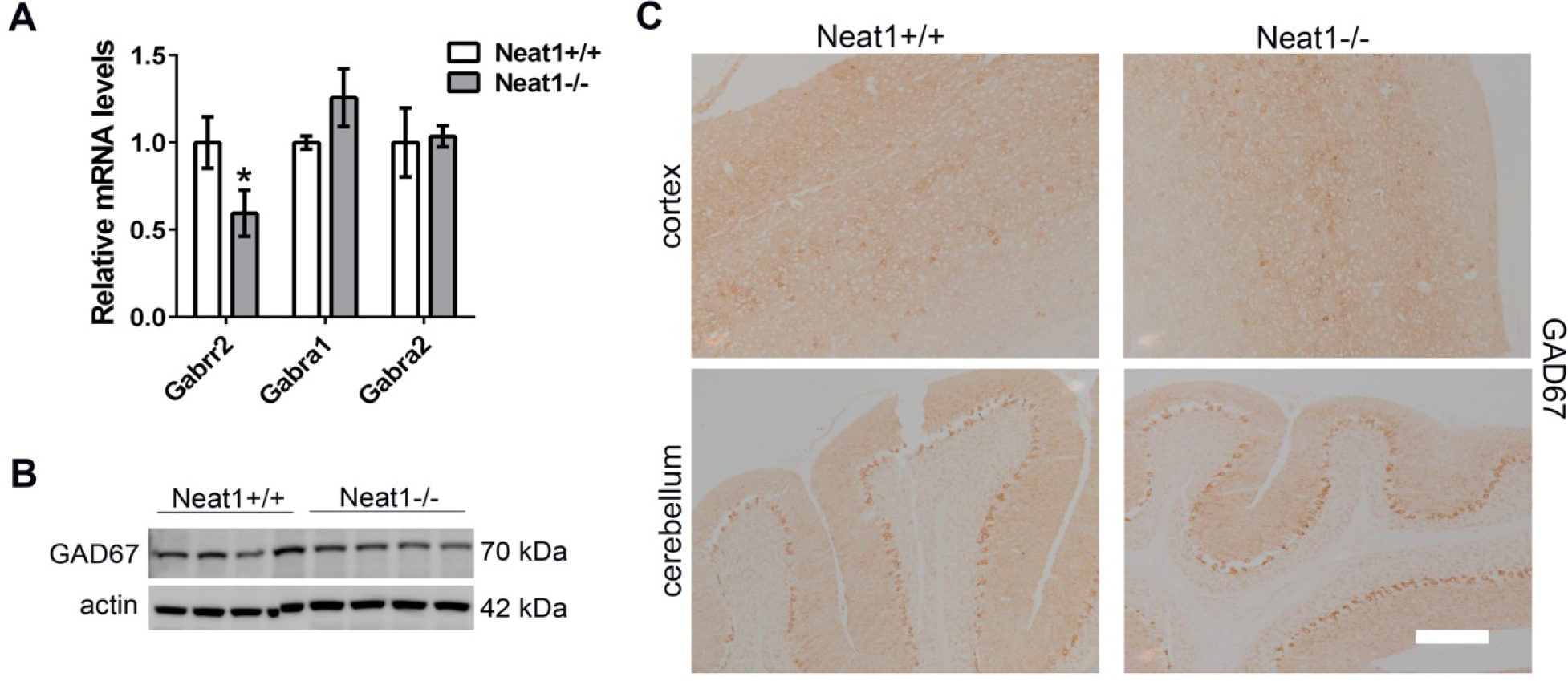
Loss of Neat1 does not affect GABAergic neurons. (**A**) Decreased expression of *Gabrr2* but not *Gabra1* or *Gabra2* in the cortex of *Neat1^-/-^* mice as revealed by qRT-PCR (Mann-Whitney *U* test; *p< 0.05, *Neat1^-/-^* n=4, *Neat1^+/+^* n=4). (**B,C**) Unaltered Gad67 levels (**B**) and distribution (**C**) in the cortex and cerebellum of *Neat1^-/-^* mice as measured by Western blot and immunohistochemistry, respectively. Representative images are shown. Scale bar, 100 µm.

**Table S1.** Proportion (%) of *Neat1^+/+^*(left) and *Neat1^-/-^* (right) mouse hippocampal neurons which demonstrated each of the different types of spontaneous action potentials (sAP type: quiet, attempting or spontaneous).

| Neat1<sup>+/+</sup> |  |  |  | Neat1<sup>-/-</sup> |  |  |  |
| --- | --- | --- | --- | --- | --- | --- | --- |
|  | Type of activity | (%) | n |  | Type of activity | (%) | n |
|  |  | 100% | 11 |  |  | 100% | 6 |
| sAP type (%) | Quiet | 18% | 2 | sAP type (%) | Quiet | 0% | 0 |
|  | Attempting | 45% | 5 |  | Attempting | 50% | 3 |
|  | Spontaneous | 36% | 4 |  | Spontaneous | 50% | 3 |

**Table S2.** Proportion (%) of *Neat1^+/+^*(left) and *Neat1^-/-^* (right) mouse hippocampal neurons which demonstrated each of the different types of induced action potentials (iAP type: none, attempting single, single, attempting train, train).

| Neat1<sup>+/+</sup> |  |  |  | Neat1<sup>-/-</sup> |  |  |  |
| --- | --- | --- | --- | --- | --- | --- | --- |
| iAP type | Total number of cells | (%) | n | iAP type | Total number of cells | (%) | n |
|  |  | 100% | 10 |  |  | 100% | 5 |
|  | None | 0% | 0 |  | None | 0% | 0 |
|  | Attempting single | 0% | 0 |  | Attempting single | 0% | 0 |
|  | Single | 30% | 3 |  | Single | 0% | 0 |
|  | Attempting train | 30% | 3 |  | Attempting train | 0% | 0 |
|  | Train | 40% | 4 |  | Train | 100% | 5 |

**Table S3.** . Comparison of passive and active parameters of *Neat1^+/+^* (left) and *Neat1^-/-^* (right) mouse hippocampal neurons. *Significantly different (p < 0.05) from *Neat1^+/+^*mouse neurons. Abbreviations: Membrane potential (Vm); input resistance (Rin), and; whole cell capacitance (Cp); I Na max and I K max are maximal values of sodium (Na^+^) and potassium (K^+^) transmembrane currents.

|  |  | Neat1<sup>+/+</sup> |  |  | Neat1<sup>-/-</sup> |  |  |
| --- | --- | --- | --- | --- | --- | --- | --- |
|  |  | Mean | SEM | n | Mean | SEM | n |
| Passive | Vm (mV) | -45.6 | 1.6 | 11 | -44.7 | 3.2 | 6 |
|  | Rin (GΩ) | 0.6 | 0.1 | 10 | 0.7 | 0.1 | 5 |
|  | Cp (pF) | 33.5 | 5.9 | 10 | 20.2 | 3.1 | 5 |
| Spike analysis | Threshold (mV) | -38.1 | 1.8 | 10 | -43.2 | 4.0 | 5 |
|  | Overshoot (mV) | 17.0 | 4.3 | 10 | 28.1 | 7.0 | 5 |
|  | After hyperpolarization (mV) | -52.1 | 2.9 | 10 | -61.4 | 5.4 | 5 |
|  | Amplitude (mV) | 69.1 | 6.2 | 10 | 89.6 | 9.9 | 5 |
|  | Depolarization rate (V/s) | 61.6 | 9.6 | 10 | 103.1* | 18.4 | 5 |
|  | Repolarization rate (V/s) | -37.9 | 8.1 | 10 | -71.6* | 14.1 | 5 |
|  | Half width (ms) | 2.8 | 0.5 | 10 | 1.7 | 0.2 | 5 |
|  | I Na max (pA/pF) | -84.5 | 18.0 | 10 | -191.3* | 61.8 | 5 |
|  | I K max (pA/pF) | 84.2 | 17.5 | 10 | 167.6 | 56.2 | 5 |

**Table S4.** List of differentially expressed genes in the cortex of *Neat1^-/-^* mice. *Available as an Excel file*.

**Table S5.** List of differentially spliced genes in the cortex of *Neat1^-/-^* mice. *Available as an Excel file*

**Video S1.** Impulsive behaviour of *Neat1^-/-^* mice upon handling.

